# Identification of novel proteins regulating lipid droplet biogenesis by reverse genetics

**DOI:** 10.1101/2022.11.24.517872

**Authors:** Md. Abdulla Al Mamun, M. Abu Reza, Md Sayeedul Islam

## Abstract

Lipid droplets (LDs) are storage organelles for neutral lipids. Our knowledge about fungal LD biogenesis is limited to budding yeast. Moreover, the regulation of LD in multinucleated filamentous fungi with considerable metabolic activity is unknown. Here, 19 LD-associated proteins were identified in *Aspergillus oryzae* using colocalization screening of a previously established Enhanced green fluorescent protein (EGFP) fusion proteins library. Following a functional screening, 12 candidates have been identified as lipid droplet regulating (LDR) proteins, the loss of which resulted in aberrant LD biogenesis. Four LDR proteins localize to LD via the putative amphipathic helices (AHs), as demonstrated with bioinformatics, targeted mutagenesis, and imaging. Further analysis revealed that LdrA with Opi1 domain is essential for cytoplasmic and nuclear LD biogenesis via this novel AH. Phylogenetic analysis demonstrated the pattern of their evolution, which was predominantly based-on gene duplication. Our study provides substantial molecular and evolutionary insights into LD biogenesis and creates a breakthrough in using reverse genetics for identifying LD-regulating proteins.

## Introduction

Lipid droplets (LD) are the ubiquitous cell organelle serving primarily as the reservoir of neutral lipids such as triacylglycerols (TG), and steryl esters (SE), which can readily be used for lipid metabolism and energy production in response to the organism’s demand (Olzmann & Carvalho, 2019; Walther et al, 2017). When the cell has an excess of fatty acids, they are converted into neutral lipids, such as TG, stored in the LDs. As energy demand increases during membrane growth, the catabolic process begins with the activation of lipolysis, which hydrolyzes the stored TG into fatty acid. Therefore, the biogenesis and breakdown of LD are tightly coupled with coordinated metabolism, and dysregulation may result in aberrant cell physiology (Welte, 2015). Furthermore, LD can also be formed inside the nucleus (Romanauska & Köhler, 2018; Sołtysik et al, 2021); therefore, its appropriate regulation is important for genomic integrity. With the recent revelation of its numerous novel functions, including host defense (Bosch et al, 2020; Monson et al, 2021) and drug delivery (Dubey et al, 2020), LD has gained further research interest.

LD consists of a neutral lipid core surrounded by a unique phospholipid monolayer, instead of the physically coupled two-leaflets conventional bilayer membrane system (Olzmann and Carvalho, 2019). During LD budding, the phospholipid monolayer originates from the cytosolic face of the endoplasmic reticulum (ER); hence a portion of ER membrane proteins such as diacylglycerol acyltransferase (DGA2) passively move to the LD surface layer (Olzmann and Carvalho, 2019; Walther et al, 2017; Choudhary et al, 2015). In contrast, a subset of cytosolic proteins binds actively to the LD via a region anticipated to contain amphipathic helix (AH) (Hinson et al, 2009; Barneda et al, 2015; Rowe et al, 2016; Prévost et al, 2018; Čopič et al, 2018; Chorlay et al, 2020; Du et al, 2020). The separation of hydrophobic and polar residues on two faces of a helix is characteristic of AH. AH enriched with large hydrophobic residues such as phenylalanine (F) exhibits a relatively strong affinity to LD surface (Prévost et al, 2018). However, weakly associated proteins are displaced from the LD surface due to protein crowding when the LD membrane shrinks during lipolysis (Kory et al, 2015).

Pezizomycotina, the largest subphylum of Ascomycota possesses research models, pathogens, and species for commercial applications (Shen et al, 2020). Pezizomycotina hyphal cells are multinucleated (Liu et al, 2009). However, the biogenesis and regulation of LD in multinucleated cells/organisms are unknown. Moreover, the number of protein-encoding genes in Pezizomycotina is roughly double that of monophyletic yeast clades Saccharomycotina and Taphrinomycotina (Nguyen et al, 2017). Numerous protein families, including enzymes for plant biomass degradation and secondary metabolism, have expanded lineage-specifically in Pezizomycotina (Arvas et al, 2007), possibly by gene duplication. These gene families are either abundant in Pezizomycotina, Pezizomycotina-specific, or lacking in Saccharomycotina (Arvas et al, 2007). Furthermore, comparative genomics with global enzyme-encoding fungal genes similarly reveals an abundance of clustered candidates related to lipid metabolism in Pezizomycotina (Wisecaver et al, 2014), implying metabolic versatility and, alternatively, the emergence of energy storage machinery to balance the potential metabolic load. However, from the viewpoint of metabolic gene evolution, LD regulation is unknown.

Here, to identify novel LD-associated proteins and patterns of LD regulation in Pezizomycotina, we employed a reverse genetics approach including colocalization screening with an EGFP-fused protein library. Following the functional screening, a group of novel candidates termed lipid droplet regulating (LDR) proteins was discovered which played an important role in maintaining LD number and size. As analyzed, at least four LDR proteins localized to the LD membrane via the putative AHs, as demonstrated by AH-disruptive mutations and subsequent imaging. Notably, the studied AH of LdrA plays an essential role in the biogenesis of cytoplasmic and nuclear LD. Using bioinformatics, we elucidated their evolutionary patterns, predominantly by gene duplication.

## Results

### Identification of novel LD-associated proteins by colocalization screening

LD biology and its associated regulation have been sufficiently studied in budding yeast. Budding yeast proteins known for LD biogenesis were compared with those of representatives from the sub-phylum Pezizomycotina, Saccharomycotina, and Taphrinomycotina of Ascomycota using BLASTp (Table S1). Despite the absence or high divergence of a portion of components such as Pln1, Ldh1, and Pgc1, many LD-regulating proteins, including key enzymes such as Pah1, Dga1, Lro1, Are2 revealed a considerable sequence homology (≤1.0e^-30^ and ≤1.0e-100 in Table S1). On the other hand, Seipin (Sei1) module/interacting proteins (such as Nem1-Spo7, Ldb16, Ldo16, and Yft2) known for ER-LD contact and crucial for regular LD biogenesis were either absent or highly diverged (>1.0e-30 in Table S1) in analyzed species. Similar divergence was also found for the OPi1 module (Opi1-Ino2-Ino4) (>1.0e-30 in Table S1) known for phospholipid regulation and LD biogenesis (Loewen et al, 2004, Romanauska & Köhler, 2018). Taking this *in silico*-based result into consideration, we hypothesize that Pezizomycotina could evolve an unidentified set of components which mechanistically support LD biogenesis.

To identify novel LD-associated proteins, we employed an EGFP-fused protein library developed by our group with a large set of uncharacterized proteins conserved in Pezizomycotina but absent or diverged in the representatives of Saccharomycotina and Taphrinomycotina (Fig 1A and Mamun et al, 2022 *preprint*). A total of 314 strains (Table S2) corresponding to the organelle-like pattern of subcellular localization were analyzed here after culturing them on the minimal medium (Fig 1A). This nutrient-limiting condition induced LD formation in *A. oryzae* (Fig 1A), similarly budding yeast (van Zutphen et al, 2014). Lipophilic fluorescent dye, Nile red (Greenspan et al, 1985) was used to stain LD in living hyphae. A total of 19 candidates were identified primarily as LD-associated with colocalization screening (12 in Fig 1B and 7 in Fig S1A). Since the EGFP-fused library was developed with a large set of candidate proteins, the identified 19 candidates were tagged with mCherry and their LD association was further validated with neutral lipid dye BODIPY 493/503 (Qiu et al, 2016) (Fig S1B and C). Twelve of the 19 proteins were functionally important for LD biogenesis and, uncharacterized of the ten were termed LDR proteins (Fig 2A–C). In addition, SppE and SppO, which participate in septal pore regulation (Mamun et al, 2022 *preprint*), were identified here to be associated with LD (Fig 1B).

**Figure 1.**
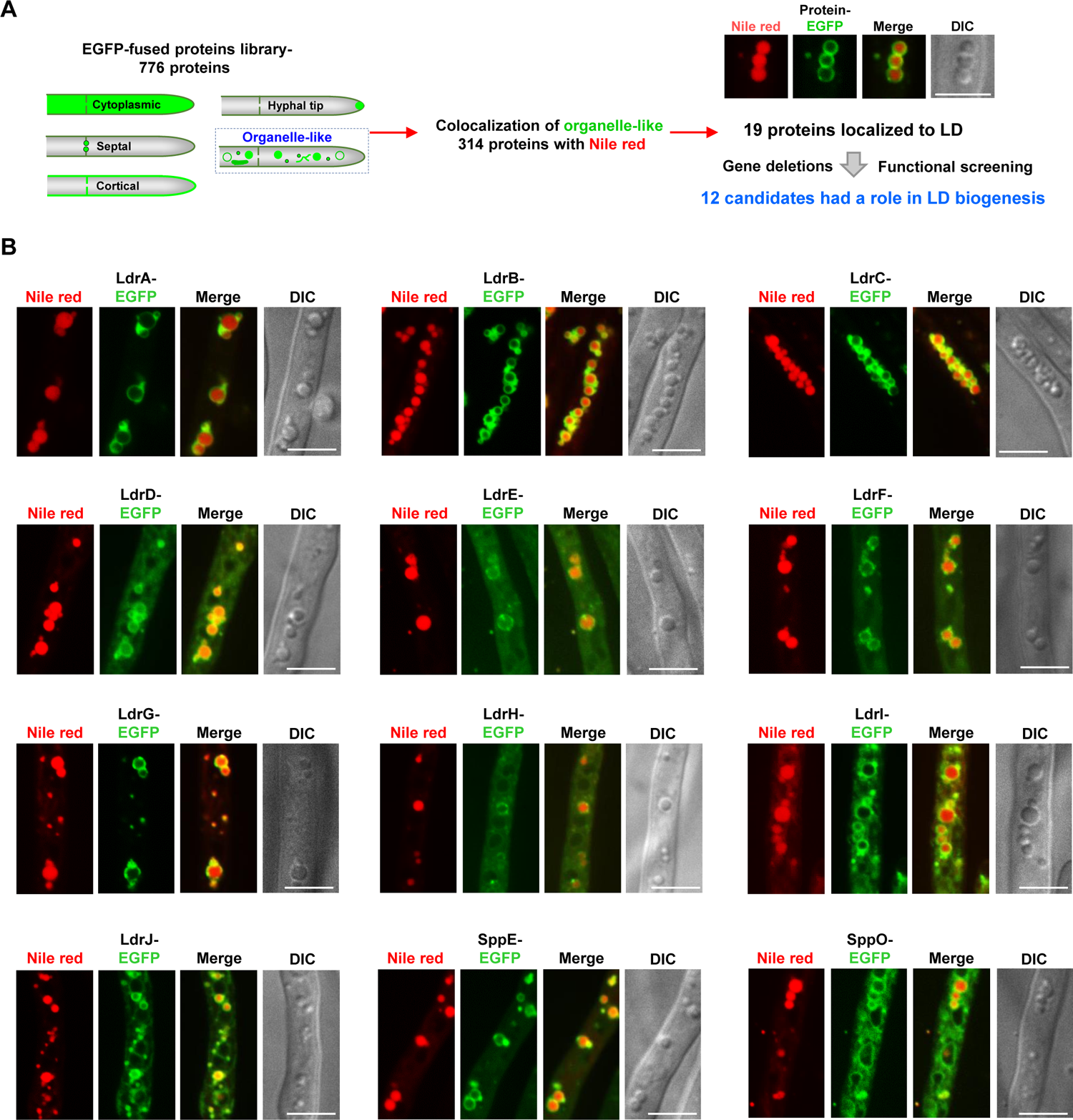
Colocalization screening identifies a set of novel proteins associated with lipid droplets. **(A)** Schematic representation of the overall strategies involved in the reverse genetic approach including colocalization and functional screening to identify novel proteins regulating LD dynamics in Pezizomycotina. **(B)** Ectopically introduced EGFP-fused LDR-, and SPP proteins, were expressed from inducible *amy*B promoters. Colocalization of EGFP-fused proteins with Nile red was observed from living hyphae cultured on the CD medium for 20 h. Fungal hyphae were stained with 10 µg/mL Nile red for 10 min. The representative microphotographs are shown. Scale bars 5µm.

**Figure 2.**
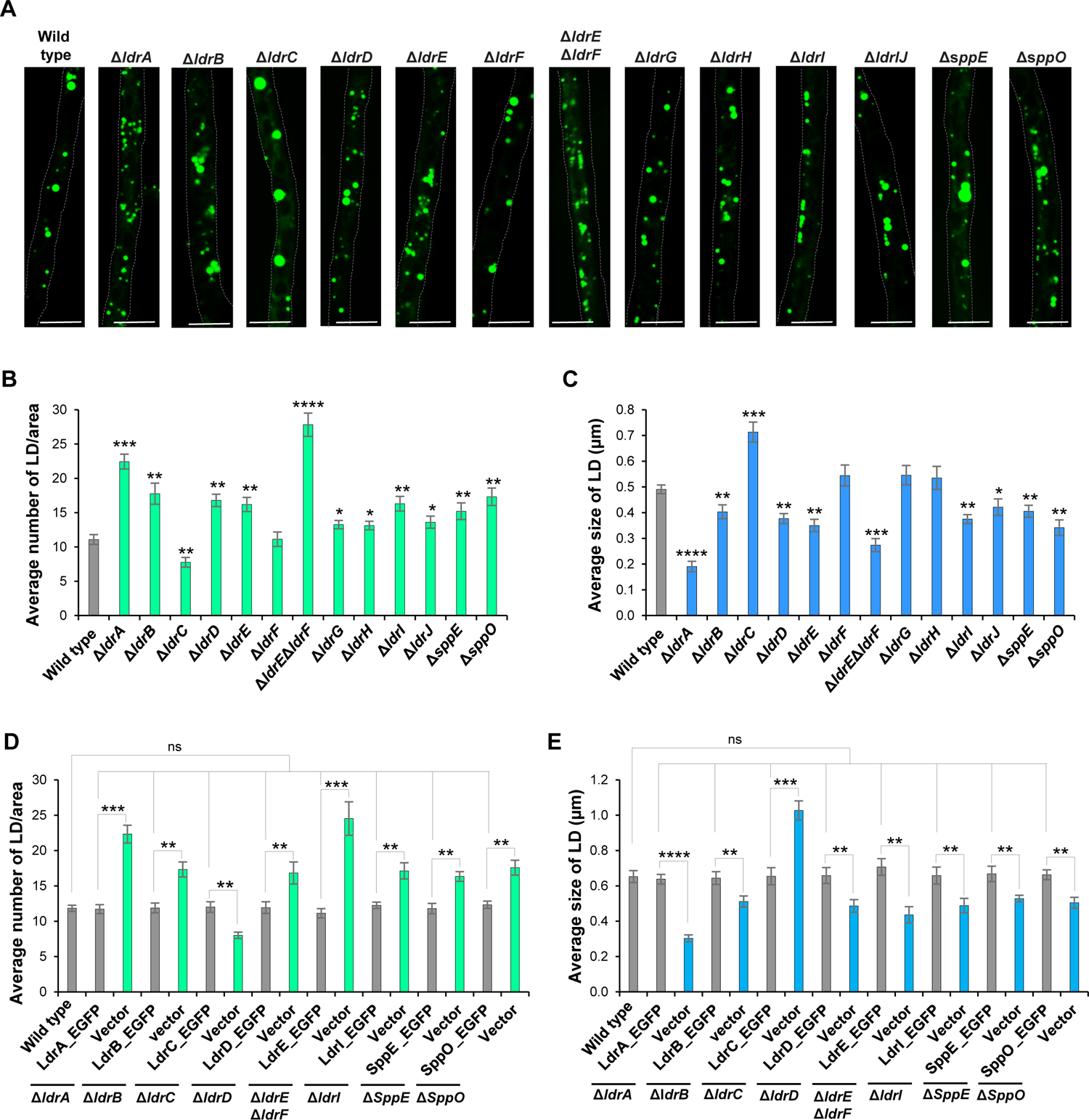
Deletions of 10 LDR–, and two SPP proteins resulted in abnormal LD number and size. LDs were quantitatively analyzed from living hyphae cultured in the minimal medium (M medium) for 20 h. Fungal hyphae were stained with 5 µM BODIPY 494/503 for 10 min. **(A)** Representative microphotographs of BODIPY-stained LD observed in wild-type and deletion strains were shown. Dotted lines represent the hyphal periphery. Scale bars 5 µm. **(B)** Quantification of LD number per hyphal area (45 µm) from ≥ 37 hyphae. **(C)** Automated quantification of the size of LD using ImageJ. n ≥ 354 LDs. **(D, E)** Deletion and double deletion strains were introduced with the respective full-length gene sequences tagged with EGFP and expressed from the corresponding endogenous promoter. Positive (wild type) and negative (vector integration) controls were developed with a similar auxotrophic background. **(D)** Quantification of LD number per hyphal area (45 µm) from ≥ 24 hyphae. **(E)** Quantification of the size of LD using ImageJ. n ≥ 313 LDs. **(B-E)** Results are represented as the mean value from three independent experiments. Statistical significance was evaluated using the two-taled student *t-test*. Error bars represent SD. ns-not significant p > 0.05, *p < 0.05, **p < 0.01, ***p < 0.001, ****p < 0.0001

Neutral lipids are synthesized between the leaflets of the ER membrane, and their subsequent growth and maturation in the forms of LD occur towards the cytosolic side (Olzmann and Carvalho, 2019; Walther et al, 2017). Next, the subcellular localization of 12 LD-associated proteins was evaluated by considering the distribution of the ER network. Markina-Iarrairaegui et al. (2013) demonstrated the ER network with the translocon component Sec63 in *Aspergillus nidulans*. Multiple LDR proteins and SppE exhibited accumulated localization as minute puncta on the ER membrane, which are likely the locations of early LD biogenesis (Fig S2). In conclusion, the current colocalization screening uncovered a group of novel candidate proteins having an association with LD.

### Role of LDR proteins in the regulation of LD number and size

Nineteen LD-associated proteins were identified using neutral lipid dyes (Fig 1 and Fig S1A-C). Next, their functions on LD biogenesis were determined after the deletion of the candidate genes. LD phenotypes in individual deletion were evaluated in living hyphae cultured on the minimal medium (Fig 2A–C). Twelve of the 19 deletion strains exhibited aberrant LD phenotypes in terms of number and size; ten of them have been characterized as LDR proteins (SppE and SppO are previously identified) (Fig 2A–C). Ten of the functionally important candidates shared the phenotype of an increase in the average number of LD but a decrease in their size (Fig 2A–C), indicating defective LD growth and maturation. Whereas, *ldrC* deletion displayed the opposite characteristics, with a drop in LD number and an increase in size (Fig 2A–C). Super-sized LD was occasionally observed in Δ*sppE* along with the predominant diminutive population (Fig 2A). Notably, among the deletion mutants, *ldrA* had the most severe LD abnormalities; over twofold increases and decreases in average number and size, respectively (Fig 2A–C). Since LdrE and LdrF as well as LdrH and LdrI were paralogues (see Fig S7B and C), double deletions were generated for further investigation of their integrated function. Despite the successful development of Δ*ldrE*Δ*ldrF*, we were unable to obtain Δ*ldrG*Δ*ldrH,* even after performing multiple round transformations in alternative combinations, which suggests that Δ*ldrG*Δ*ldrH* might have a deleterious effect on the organism. Importantly, Δ*ldrE*Δ*ldrF* showed severe phenotypes in terms of LD number and size compared with either single deletion (Fig 2A-C).

Expression of EGFP-fused individual gene sequence from the relevant endogenous promoter restored defective LD features (Fig 2D and E) observed in the corresponding deletion, demonstrating the functional relevance of the candidate proteins in LD biogenesis and the validity of the EGFP-fused constructs. Nonetheless, some EGFP-fused candidates were detected on the LD surface, but others were not, probably because of the weak EGFP signal under native expression (Fig S1D). Finally, with functional screening, we identified LDR proteins that played an important role in maintaining LD number and size.

### LD**–**localized proteins associate with nascent LD-marker proteins

Diacylglycerol acyltransferase (Dga1) catalyzing the terminal step of TG biosynthesis concentrates at the subdomains of the ER membrane, which is known to be the site of early LD biogenesis (Joshi et al, 2018; Choudhary et al, 2020; Jacquier et al, 2011). Erg6, a component of the ergosterol biosynthesis pathway, is also being employed as a marker of early-stage LD (Joshi et al, 2018; Choudhary et al, 2020; Jacquier et al, 2011). Consistently, AoDga1- and AoErg6-EGFP colocalized predominantly with Nile red-stained LD (Fig S3A). Several LD-associated proteins exhibited accumulated localization on the ER membrane (Fig S2). Next, we determined if they were associated with AoDga1 and AoErg6. Among the analyzed proteins, LdrA-, LdrB-, LdrG-, and SppE-EGFP were largely associated with both AoErg6- and AoDga1-mCherry or *vice versa*, either as puncta or as the rim of LD (Fig 3A– D), similar to their regular localization to the LD (Fig 1B). However, LdrC-, LdrI-, LdrF-, and LdrD-EGFP exhibited a partial displacement from LD upon co-expression (Fig S3B–E), compared with their regular pattern of localization (Fig 1B). Notably, AoErg6-mCherry also partially lost its exclusive association with LD when co-expressed with LdrB-, LdrC-, and LdrI-EGFP (Fig 3B and Fig S2B and C). Interestingly, AoErg6-mCherry regained its natural punctate or rim-like localization (Fig S3B) when co-expressed with the less effective variant LdrC(V175N) (See Fig 6C), implying a possible competition of their association to the surface of LD, as previously reported (Kory et al, 2015). Finally, several of the examined LD-regulating proteins were predominantly associated with AoErg6 and AoDga1, frequently as tiny puncta (arrowheads in Fig 3A-D), suggesting their participation in nascent LD biogenesis.

**Figure 3.**
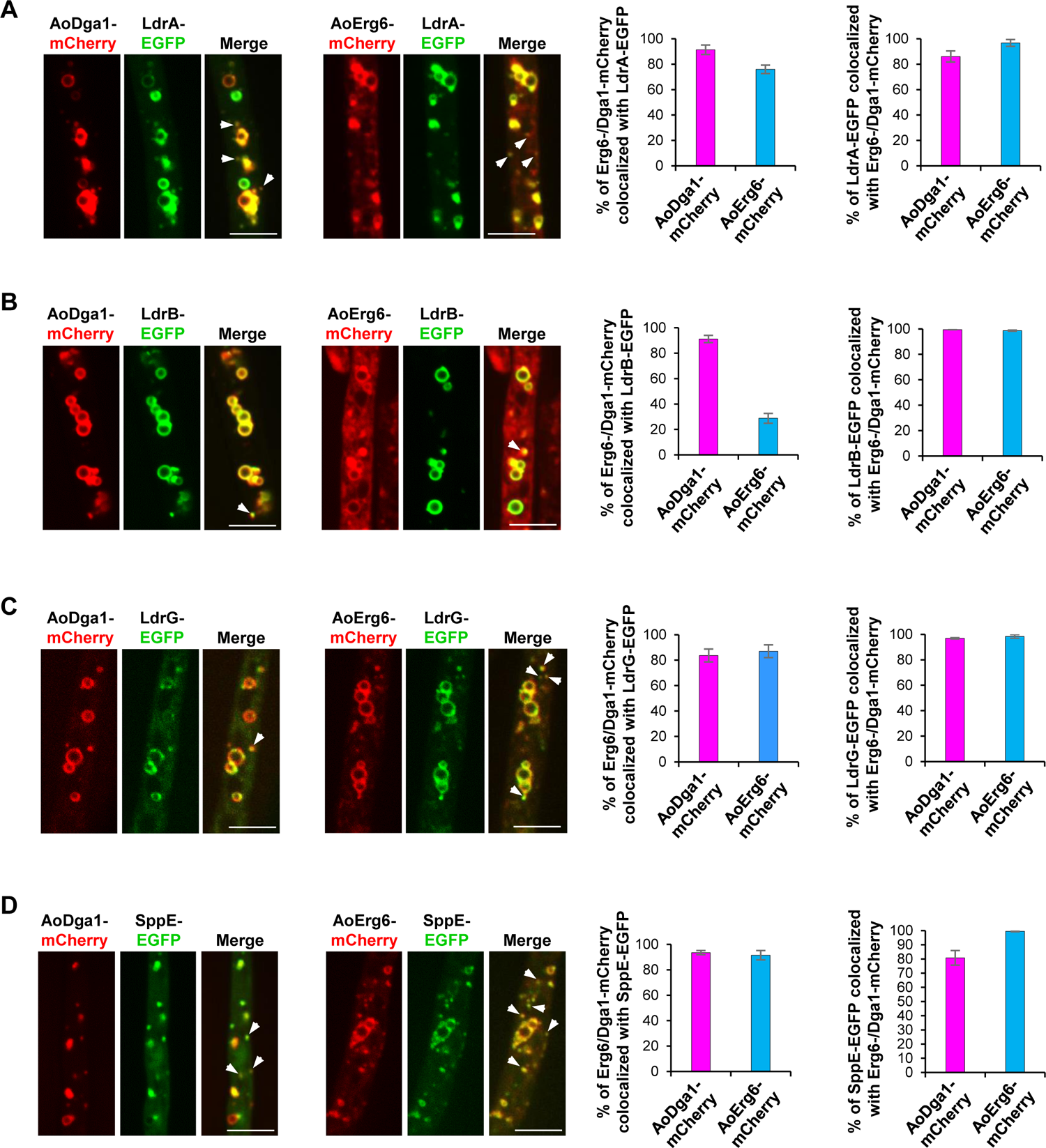
Colocalization of four LD-associated proteins with nascent LD marker Erg6 and Dga1. AoErg6- and AoDga1-mCherry under the control of *amyB* promoter were ectopically introduced individually into the wild-type. The plasmids bearing EGFP-tagged genes corresponding to LDR proteins were further ectopically introduced into the resultant strains expressing mCherry labeled AoErg6 and AoDga1. Strains were cultured on the CD medium. Representative microphotographs were shown. Arrowheads represent the colocalization points at small-sized puncta. Scale bars 5µm. The percentage of AoErg6- and AoDga1-mCherry associated with EGFP-tagged LDR proteins or *vice versa* were calculated using ImageJ. Data are presented as mean ± SD, n = 12 hyphal area from three independent experiment. **(A)**, **(B)**, **(C)**, and **(D)** colocalization analysis of AoErg6- and AoDga1-mCherry with LdrA-, LdrB-, LdrG-, and SppE-EGFP, respectively.

### LdrA binds to and regulates LD biogenesis via a novel AH

During LD maturation, membrane curvature causes phospholipid (PL) packing defects (Olzmann and Carvalho, 2019). A subset of AH-containing proteins utilizes this property to directly binds from the cytosol to the LD membrane, hence promoting membrane curvature and subsequent LD growth (Bigay et al, 2012; Prévost et al, 2018; Chorlay et al, 2020). Several LDR proteins had a rim-like localization to the LD membrane (Fig 1B). Next, we determined whether LDR proteins particularly LdrA possess the sequence characteristic predicted to be the AHs that provide their binding to LD. To predict the potential AH, total α-helices were comprehensively retrieved using Phyre2 (Kelley et al, 2015) and then analyzed with HELIQUEST (Gautier et al, 2008). The helix was primarily selected if hydrophobic and polar residues showed segregation with the hydrophobic moment around 0.35. The predicted AHs were further evaluated with AlphaFold2 (Mirdita et al, 2022).

With this comprehensive analysis, LdrA was predicted to contain a region at the C-terminus having the potential to be an AH with a hydrophobic moment of 0.34 (Fig 4A). Importantly, the positions of hydrophobic and polar residues were conserved across the fungal representatives (Fig 4B) implying evolutionary conservation of the AH. To disrupt the putative AH, hydrophobic residues were substituted with polar asparagine (N), which maximizes the reduction of the overall hydrophobicity without altering its charge (Fig 4C). The size and expression of the truncated and mutated variants were evaluated using western blotting (Fig S4A). The variants were analyzed for any alteration of their subcellular localization with Nile red. LdrA variant lacking AH, LdrA(ΔAH) was completely nuclear (Fig 4C). A progressive decline in hydrophobicity led to the variants translocating into the nucleus (Fig S4B–D and Table 1), marked with the component of nuclear pore complex (NPC), Nup84 (Liu et al, 2009). In contrast, except for the AH dead mutant, LdrA(V420N/V416N), the variants were still associated with LD (Fig 4C and Fig S4E, F). This inconsistent result can be explained by a markedly increased expression (protein) of those variants persisting functional AH (Fig S4A and Table 1). Moreover, replacing hydrophobic residues with similar properties such as valine to leucine (V420L) did not affect their regular subcellular localization to the LD membrane (Fig 4C and Fig S5A, B).

**Figure 4.**
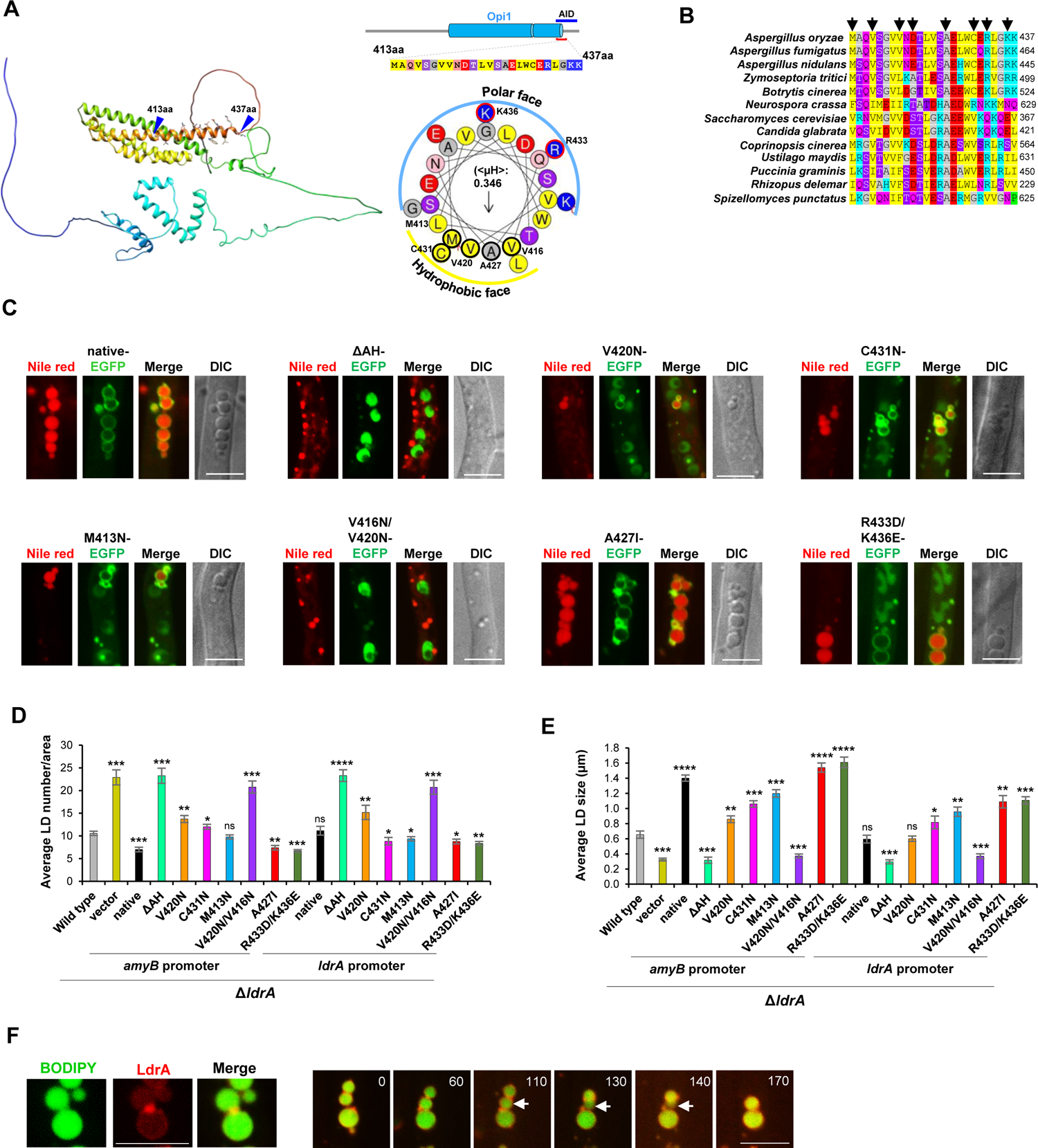
LdrA binds to and regulates LD biogenesis via the insertion of a putative amphipathic helix. **(A)** Ribbon diagram of the 3D structure of LdrA were predicted using AlfaFold2. N-terminus is blue, C-terminus is red (rainbow orientation). Putative AH at the C-terminus is presented with side chain. Helical wheel representation of predicted AH using HeliQuest. Amino acid properties are indicated by colors. Mutated residues were highlighted with bold borders. **(B)** Multiple sequence alignment of AH using ClustalW. The sequences were retrieved from representative fungal orthologues. Arrows denote the residues that were substituted for AH-disruption. **(C)** The constructs designed to disrupt the amphipathic nature of AH, expressed from the *amyB* promoter were introduced into Δ*ldrA.* Live-cell imaging was performed from the LdrA variants cultured on the CD medium. Representative images were shown. Scale bars 5 µm. **(D)** and **(E)** Quantification of LD number per hyphal area (45µm) and LD size from LdrA variants (untagged), respectively. LD was visualized with BODIPY 493/503. The bar diagrams are represented as the mean value from three independent experiments. **(D)** n ≥ 27 hyphae. **(E)** n ≥ 135 LDs. Statistical significance was assessed using two-tailed student *t-test*. Error bars represent SD. ns-not significant p > 0.05, *p < 0.05, **p < 0.01, ***p < 0.001, ****p < 0.0001. **(F)** LdrA accumulated at the LD**–**LD fusion site. Strain expressing LdrA-mCherry was stained with BODIPY. Time course observation of LD fusion (right). Arrowheads indicate ongoing fused-LD. time= min.

**Table 1:**
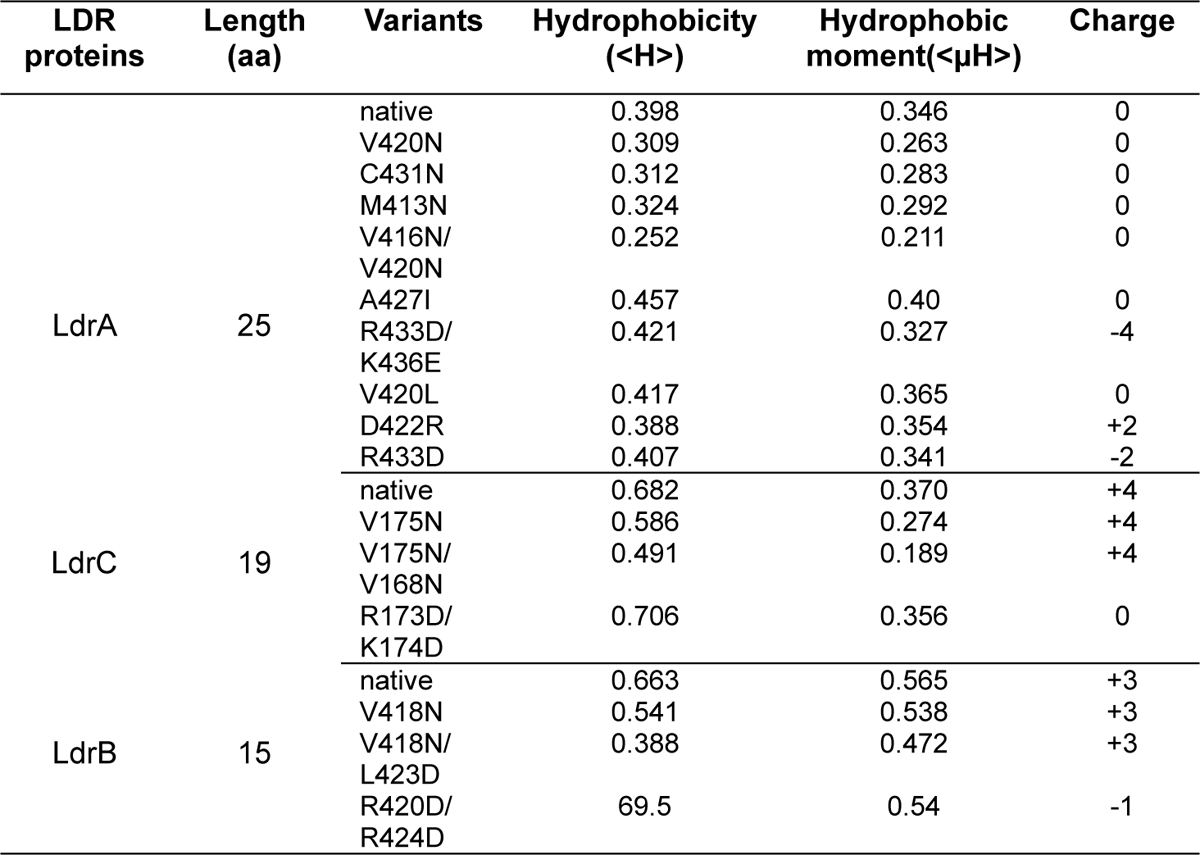
Amphipathicity comparison among the mutated variants of three LDR proteins.

LdrA possesses 0 net charge. Charge swap mutants were generated with the substitution of positively charged residues such as arginine (R) to negatively charged aspartic acid (D) to better understand their importance in LD association. Gradual enrichment of the negative charge from 0 to −4 led the variants to be partially displaced from the LD to the nucleus (Fig 4C and Fig S4B–D). At the same time, variant LdrA(D422R) with positive charge +2 did not affect the regular subcellular localization on LD (Fig S5A, B and Table 1). Taken together, our data demonstrated that the amphipathic features including hydrophobicity and positive charge played a crucial role in the association of LdrA to the LD surface.

We then investigated if the AH has a role in LD biogenesis. LdrA variants lacking AH, LdrA(ΔAH), and AH dead mutant, LdrA(V420N/V416N) produced many tiny LDs, similar to the deletion of *ldrA* (Fig 4D and E). The basal number of LD increased in the order of decreasing hydrophobicity LdrA(A427I)< LdrA (V420L)< wild type < LdrA (M413N)< LdrA (C431N)< LdrA (V420N)< LdrA (V420N/V416N), but LD size followed the order of increasing hydrophobicity (Fig 4D, E and Fig S5C, D and Table 1), demonstrating an apparent relationship between LD phenotype and hydrophobicity. Charge swap mutant LdrA(R433D/K436E), on the other hand, demonstrated an increasing LD size even the level above the wild type, presumably due to higher protein level and/or hydrophobicity (Fig 4D, E and Fig S4A and Table 1). LD fusion could be one of the possible explanations reasoned for the generation of supersized LD in variants with potential hydrophobicity, such as LdrA(A427I), because LdrA-mCherry enriched at the LD–LD contact site (Fig 4F) perhaps facilitates LD fusion. Taken together, the amphipathic features such as hydrophobicity and charge of putative AH of LdrA played a crucial role in regulating normal LD biogenesis. Furthermore, variants with impaired LD biogenesis, such as LdrA(ΔAH) showed a defect in asexual spore development (Fig S5F). Whereas variants with high LD load, such as LdrA(A427I), showed a partial decrease in colony development (Fig S5F), implying a negative relationship between lipid storage and hyphal growth.

### LdrA induces the biogenesis of nuclear LD

Budding yeast OPi1 binds to and induces nuclear LD (nLD) after losing its association with the ER (Romanauska & Köhler, 2018). Next, we asked whether LdrA is associated with nLD. The nuclear structure was visualized with the nuclear localizing signal (NLS) derived from the transcription factor StuA (Suelmann et al, 1997). EGFP-fused LdrA was co-expressed with mCherry-labeled NLS. Contrary to budding yeast Opi1, LdrA was regularly and more efficiently associated with nLD compared with cytoplasmic LD (cLD) (Fig 5A). We, next asked whether LdrA is involved in the regulation of nLD biogenesis and whether AH has any role. Untagged variants of LdrA from both inducible *amyB* and endogenous promoters were co-expressed with mCherry-labeled NLS. The current study eliminated apical compartments from observation which are known to be the active zone for nuclear division (Kaminskyj et al, 1998). BODIPY–positive foci that were identified as nLD were exhibited as a shadow of their shape on the nucleus (Fig 5B). In the wild-type, approximately 7% of the nuclei bore nLD, while this ratio increased to 20% in the variant that overexpressed wildtype LdrA (denoted as native) (Fig 5B and C). Surprisingly, 40% of nuclei were positive for nLD in LdrA(M413N) (Fig 5B and C), the variant showing dramatically increased expression (protein) and persisting maximum hydrophobicity among AH-disruptive mutants (Fig S4A and Table 1). On the other hand, the percentage of nLD was noticeably decreased in the truncated variant lacking AH LdrA(ΔAH), and AH dead mutant LdrA(V420N/V416N), with a nearly similar level observed in the deletion of *ldrA* (Fig 5B and C). However, the variants such as LdrA(A427I) and LdrA(V420L), generated by replacing hydrophobic residues with similar properties followed a similar trend to increase the nLD ratio (Fig 5B, C and Fig S5E), which collectively suggested that LdrA played an important role in nLD biogenesis via the AH.

**Figure 5.**
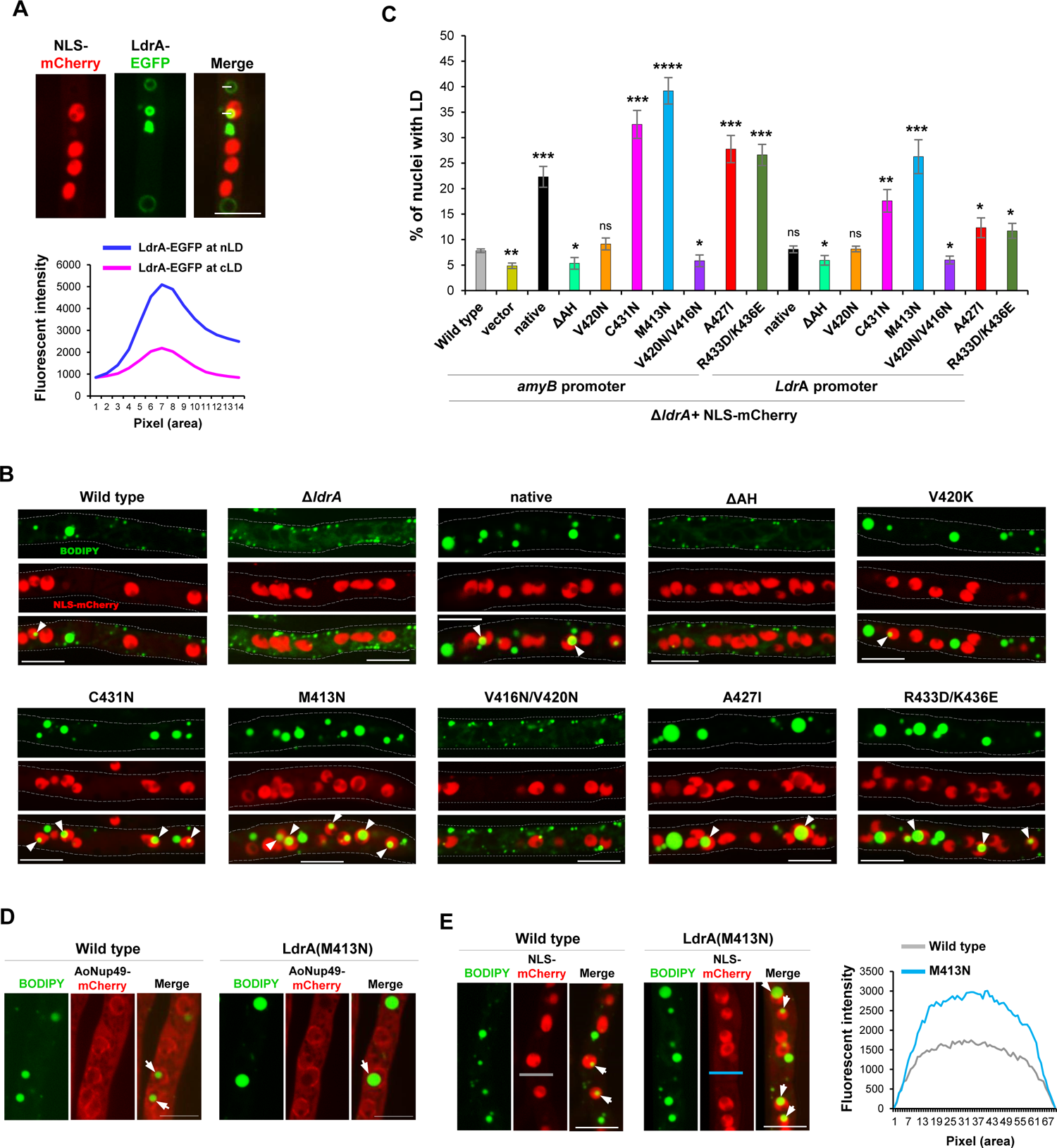
LdrA induces nuclear LD biogenesis. **(A)** Colocalization analysis of LdrA-EGFP and NLS(AoStuA)-mCherry revealed predominant binding of LdrA to nuclear LD over cytoplasmic LD. Y axis of the line graph presents the mean florescent intensity of LdrA-EGFP on LD membrane. n = 25 LDs. **(B)** Δ*ldrA+*NLS-mCherry complemented with LdrA variants, expressed from heterologous *amyB* promoter. LDs and nuclei were visualized using BODIPY and NLS-mCherry, respectively. Representative images are shown. Arrowheads indicate nLD. Dotted lines represent the hyphal periphery. Scale bars 5 µm. **(C)** LdrA variants were individually expressed from its endogenous promoter and *amyB* promoter. The bar diagrams showing the percentage of nLD are presented as the mean value from three independent experiments. n ≥ 216 nuclei. Statistical significance was assessed using the two-taled student *t-test*. Error bars represent SD. ns-not significant p > 0.05, *p < 0.05, **p < 0.01, ***p < 0.001, ****p < 0.0001. **(D)** Disruption of nuclear periphery by super-sized nLD. Nuclear periphery was visualized with AoNup49-mCherry. Arrows represents nLD. **(E)** Mean fluorescent intensity of NLS-mCherry quantified from the cytoplasmic pool of wild type and LdrA(M413A) using ImageJ. n= 20 hyphae.

Furthermore, supersized nLD may lead to nuclear disintegration (Fig 5B). Nup49, a component of the NPC (Liu et al, 2009), fused with mCherry, was introduced in the wild-type, and LdrA(M413N) variant to clearly define the nuclear periphery and the presumed nuclear disintegration (Fig 5D). As expected, the nuclear periphery was disintegrated due to the supersized nLDs in LdrA(M413N) (Fig 5D). Following that, we looked into the degree of nuclear disruption in LdrA(M413N). NLS-mCherry which is entirely nuclear was used to visualize nuclei. The mCherry intensity captured from the cytoplasmic pool was found greatly elevated in LdrA(M413N) compared to that in the wild-type control (Fig 5E), which indicated that the leaking nucleus led to the escape of the NLS-mCherry in the cytoplasm, and alternatively nuclear disintegration.

### Role of predicted AHs of LdrC, LdrB, and LdrG in their association with LD

The putative AH of LdrA is crucial for its association with LD. Using the aforementioned comprehensive screening (Fig 4A), we next determined whether the other LDR proteins possess regions that have the potential to be the AHs. A middle region comprising 19 residues of LdrC was predicted as AH, which possesses a comparatively long hydrophobic face (Fig 6A and B, Table 1). The ribbon diagram generated with AlphaFold2 confirmed the helix which could locate at the surface of LdrC (Fig 6A). Multiple sequence alignment revealed that the putative AH is conserved among the fungal representatives (Fig S6A). To evaluate the importance of predicted AH in the association of LdrC with LD, a truncated variant lacking the AH was generated. In addition, AH-disruptive variants were created with the substitution of hydrophobic residues with polar asparagine (N). Variants lacking the AH, LdrC(ΔAH) led to its complete dissociation from the LD surface (Fig 6C). Additionally, the LD affinity of the substituted variants decreased dramatically with the progressive decline of hydrophobicity in the following order: wild-type > LdrC(V175N)> LdrC(V175N/V168N) (Fig 6C and Fig S6D and Table 1). Additionally, charge swap mutants were generated by replacing positively charged residues with negatively charged aspartic acid (D), and these modifications had a negative effect on the association of the variants to LD (Fig 6C and Fig S6D and Table 1).

**Figure 6.**
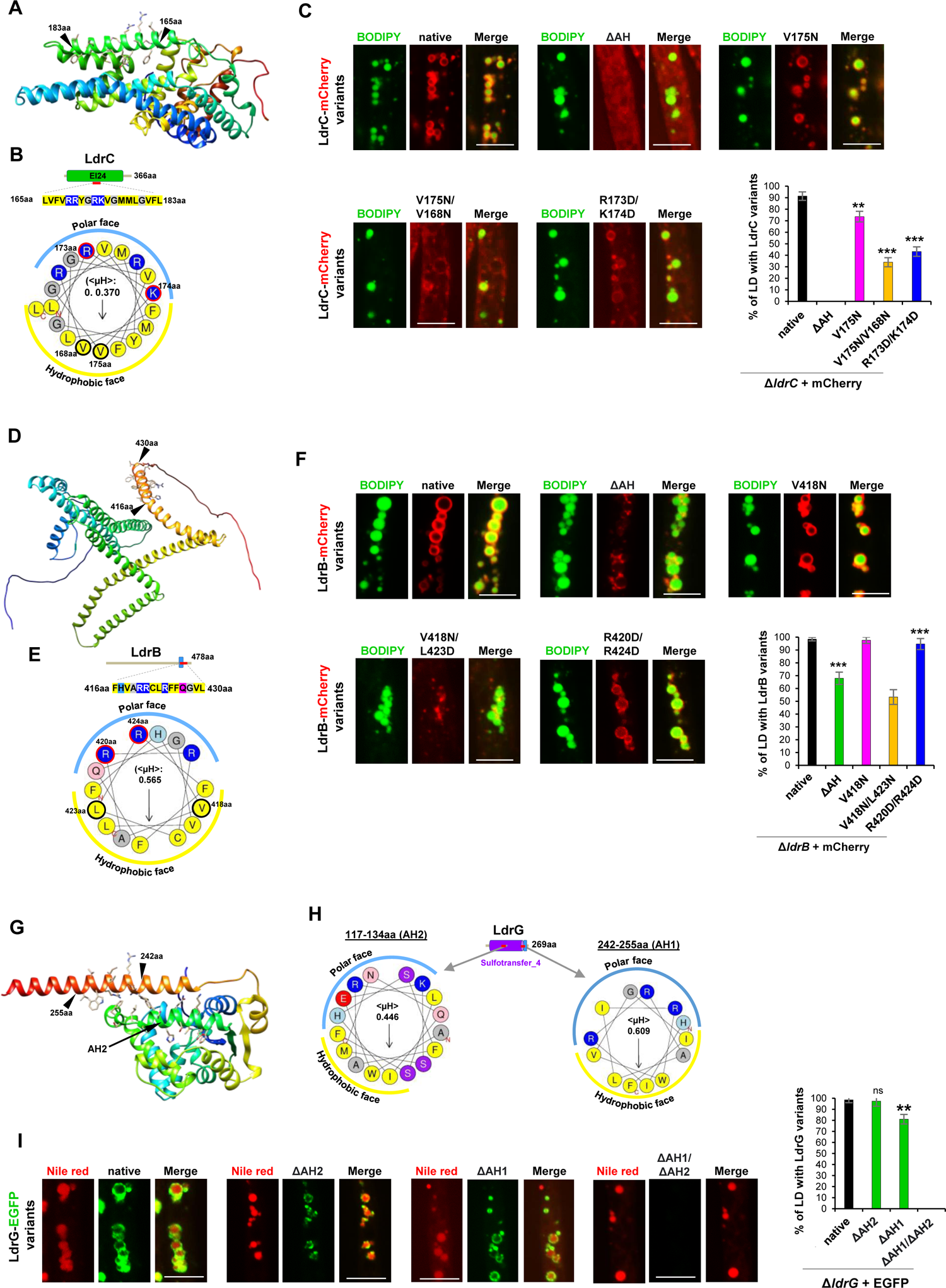
LdrC, LdrB, and LdrG associate with LD via the predicted AHs. Ribbon diagram of the 3D structure of LdrC. **(A)**, LdrB **(D)**, and LdrG **(G)** were predicted using AlfaFold2 and represented with Chimera. N-terminus is blue, C-terminus is red (rainbow orientation). Putative AHs are presented with side chains. Helical wheel representations of the predicted AHs of LdrC **(B)**, LdrB **(E)**, and LdrG **(H)** using HeliQuest server. The properties of amino acid are indicated by colors. Mutated residues were highlighted with bold borders. Hydrophobic moment = (<µH>). The constructs designed to disrupt the amphipathic nature of corresponding AH of LdrC **(C)**, LdrB **(F)**, and LdrG **(I)**. Δ*ldrC,* Δ*ldrB,* and Δ*ldrG* were introduced with the corresponding wild type and AH-disruptive mutational constructs. Hyphae cultured on CD medium were stained with BODIPY 493/503 (C, F) and Nile red (I). Scale bars 5 µm. Percentage of LD positive with LdrC, LdrB, and LdrG variants were counted with ImageJ. **(C, F, I)** Experiments were performed in triplicates. n ≥ 95 LDs. Statistical significance was assessed using two-tailed student *t-test*. Error bars represent SD. ns-not significant p > 0.05, *p < 0.05, **p < 0.01, ***p < 0.001, ****p < 0.0001.

A C-terminal region consisting of 15aa (416-430aa) of LdrB was predicted as AH (Fig 6D and E). However, this AH was conserved solely within Aspergillus species (Fig S6B). The truncated variant losing this predicted AH resulted in a marked abolishment of its uniform rim-like localization but still showed a puncta-like attachment to the LD membrane (Fig 6F). Then AH-disruptive mutants were generated for further confirmation. LD association activity of the variants was markedly decreased with the progressive decline of hydrophobicity and also positive charge (Fig 6F and Fig S6E and Table 1).

On the other hand, two AHs with individual hydrophobic moments of 0.609 and 0.446 were predicted in LdrG (Fig 6G and H). The extended organization of AH1 could facilitate the binding of LdrG to the LD membrane (Fig 6G). The positions of the polar and hydrophobic residues were mostly conserved in paralogue, LdrH, but partially conserved in the fungal orthologous (Fig S6C). The variants losing either AH revealed minor or partial effect, whereas that losing both showed the abolishment of the EGFP signal (Fig 6I), implying that unbound LdrG is unstable in the cytosol.

### The regulation of LD biogenesis from LdrA and LdrC

Because LdrA and LdrC showed opposite phenotypes in terms of LD number and size under both deleted and overexpressed conditions (Figs. 2A–C and 4D, E and Fig S2), an extended investigation was conducted to elucidate the mode of LD regulation from their relation. Firstly, double deletion of the *ldrAldrC* was generated to evaluate their integrated function in LD regulation. Interestingly, double deletion of *ldrAldrC* resulted in an LD feature that partially and completely rescued the severe LD defective phenotypes found in Δ*ldrA and ΔldrC,* respectively (Fig 7A). Additionally, the insertion of LdrA (overexpressed from *amyB* promoter) in Δ*ldrC*Δ*ldrA* led to a further increase in basal LD size (Fig 7A), suggesting that in the absence of LdrC, LdrA function could be induced.

To further investigate the mode of LD regulation, LdrA-EGFP, and LdrC-mCherry were co-expressed. Co-expression resulted in a marked removal of LdrA-EGFP from the LD surface (Fig 7B). Alternatively, the LD membrane was predominantly bound by LdrC-mCherry (Fig 7C). Interestingly, co-expression of LdrA with the less efficient variant of LdrC, LdrC(V175N/V1678N) (Fig 6C) resulted in an apparent improvement in the colocalization efficiency (Fig 7B), which collectively suggests that wild-type LdrC has a predominant association with LD, which might inhibit or decrease the accessibility of LdrA to the LD.

**Figure 7.**
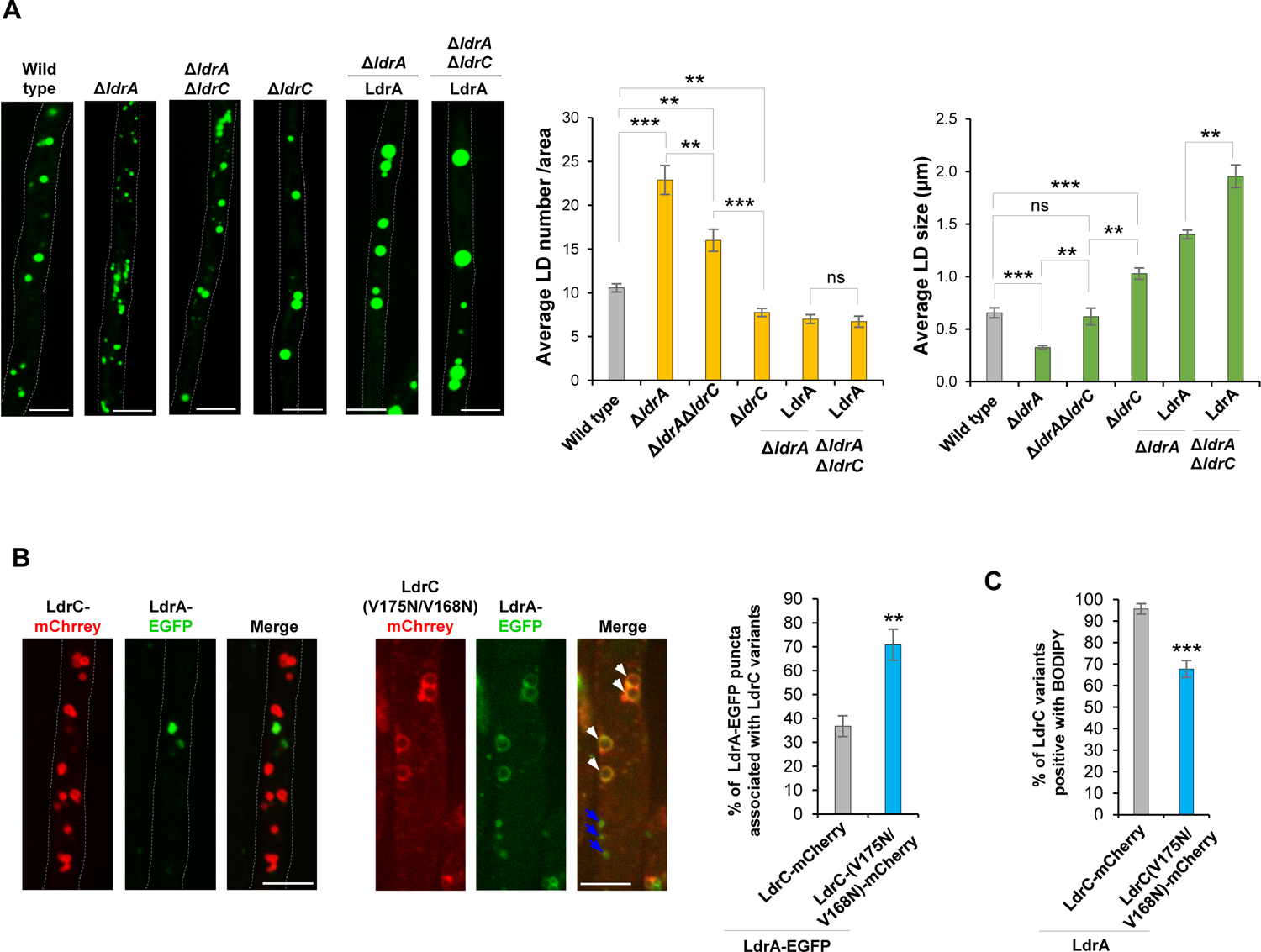
LD regulation from LdrC and LdrA. **(A)** LdrA was overexpressed into Δ*ldrA* and Δ*ldrC*Δ*ldrA* from inducible *amyB* promoter. LD number and size were analyzed using ImageJ. For LD number n ≥ 21 hyphal areas and LD size n ≥ 145 LDs. Representative microphotographs of BODIPY-stained LD are shown. **(B)** Co-expression of LdrA-EGFP and LdrC variants labelled with mCherry from *amyB* promoter. Live-cell imaging was performed from the strains cultured on the CD medium with optimized carbon concentration. White and blue arrowheads represent the colocalized and non-colocalized LdrA-EGFP, respectively. The percentage of LdrA-EGFP puncta associated with LdrC-mCherry variants was calculated. n ≥ 141 LdrA-EGFP puncta. **(C)** Co-expression of LdrA (untagged) and mCherry-labelled LdrC variants. Hyphae were stained with BODIPY. n = 68 LDs. **(A-C)** Results are represented as the mean values from three independent experiments. Statistical significance was assessed using two-tailed student *t-test*. Error bars represent SD. ns-not significant p > 0.05, *p < 0.05, **p < 0.01, ***p < 0.001. Scale bars 5 µm.

### Phylogenetic distribution and evolution of LDR proteins

We analyzed the phylogenetic distribution and the degree of divergence of LDR proteins with respect to broad data of protein sequences from representative species covering all major fungal phyla and subphyla (Ahrendt et al, 2018) using BLASTp (Fig 8 and Table S3A-C). Except for LdrA and LdrB, our results showed that LDR proteins evolved predominantly by gene duplication.

**Figure 8.**
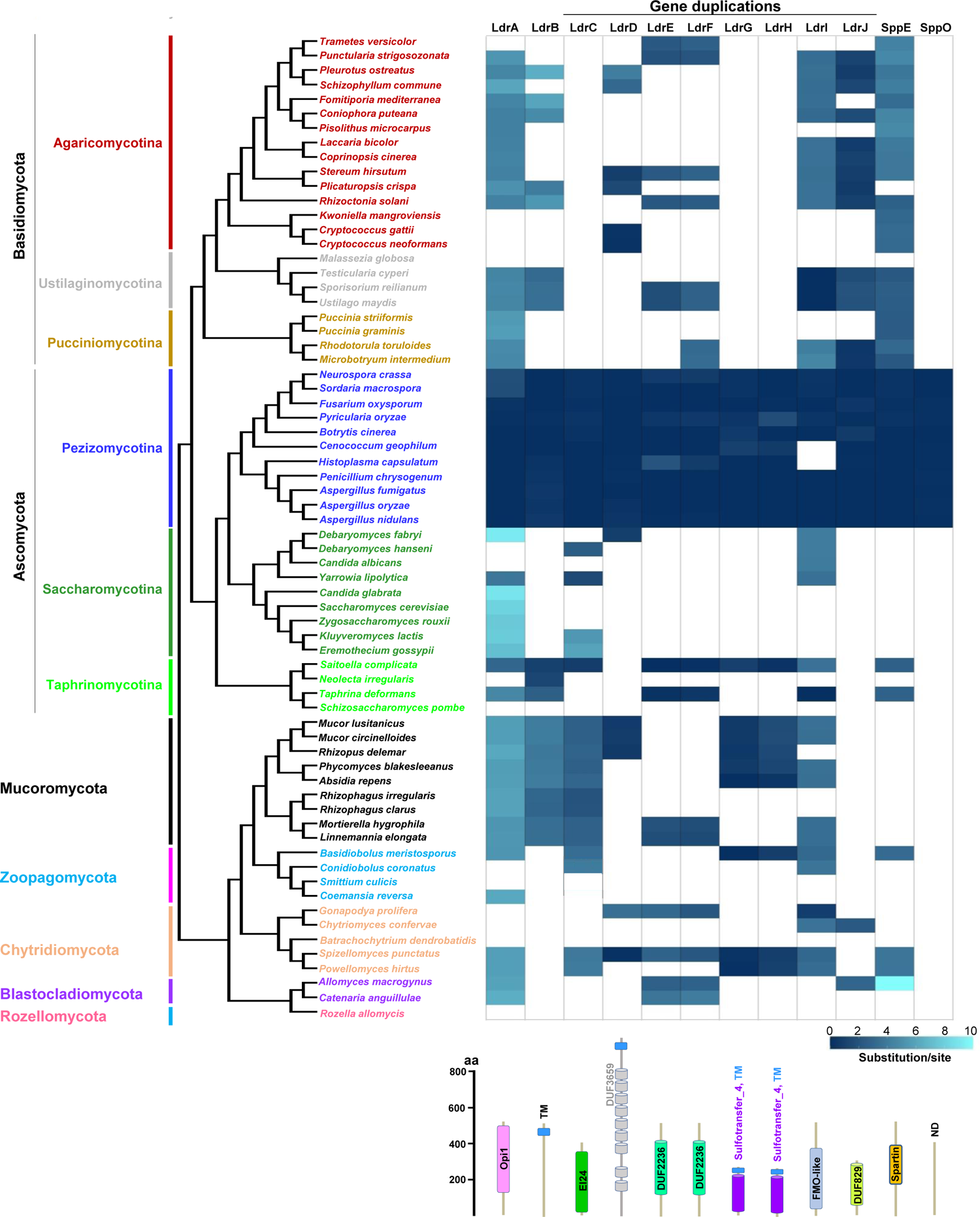
Conservation and divergence of LDR proteins across fungal phyla Phylogeny shows representative species from the major phyla or sub-phyla. Filled and empty boxes denote the presence and absence of LDR proteins, respectively, according to the specified e-value cutoff (1E+10) of BLASTp (Table S3A). The scale besides the cartoons denoting protein domains at the bottom, represents the number of amino acids. (ND, no domain). Substitution rate analysis of LDR and two SPP proteins in the fungal phylum or sub-phylum from Pezizomycotina root. Amino acid substitutions per site are shown in blue as a heat map (Table S3C). A lighter shade indicates greater divergence from the Pezizomycotina.

LdrA possessing the OPi1 domain (PF08618) (Loewen et al, 2004) showed divergence from the representatives of Saccharomycotina including budding yeasts Opi1 (Fig 8). LdrB containing transmembrane domain exited in Pezizomycotina, Mucoromycota, and a part of Basidiomycetes, but absent in Saccharomycetes (Fig 8).

Notably, a total of eight LDR proteins LdrC, LdrD, LdrE, LdrF, LdrG, LdrH, LdrI, and LdrJ evolved by gene duplications. LdrC contains EI24 domain (PF07264) (Xu et al, 2022). Although basidiomycetes and several yeast species lack–, representatives from early diverging phylum possess the orthologs (Fig 8 and Fig S7A). LdrE and LdrF contain the DUF2236 domain (PF09995). LdrG and LdrH possess the sulfotransfer-4 domain (PF17784) (Negishi et al, 2001). Notably, LdrE and LdrF, as well as LdrG and LdrH are paralogues, respectively (Fig S7B and C), indicating gene duplication. Except for Saccharomycetes, LdrE, as well as LdrF, dispersedly existed across the fungal phyla (Fig 8). On the other hand, LdrG and LdrH were found abundant in early diverging Mucoromycota and Chytridiomycota; they are absent in Basidiomycetes and Saccharomycotina (Fig 8 and Fig S7C). However, except for ascomycete yeasts, LdrJ containing the DUF829 domain (PF05705) predominantly existed in Dikarya (Fig 8 and Fig S7D). Interestingly human has protein with similar domain organizations and this possibly is orthologues as assayed by reciprocal BLAST. LdrD containing DUF3659 domain (PF12396) showed scattered existence across the fungal representatives (Fig 8). It has paralogs (Fig S7E), indicating gene duplication. However, the subcellular localization of the paralogs was independent of LD (Fig S7F). Remarkably, several orthologues showed N-terminal truncation. Multiple sequence alignment revealed a higher sequence homology with the C-terminal region of LdrD (Fig S7G), implying the protein structural evolution among the fungal orthologues. LdrI, which has an FMO-like domain (PF00743) (Lawton et al, 19940), forms a distinct clade consisting of Pezizomycotina and Ustilaginomycotina, as well as a few species of Chytridiomycota and Taphrinomycotina (Fig 8 and Fig S7H).

Finally, our data showed the existence of multiple proteins with almost similar domain organizations corresponding to eight of the LDR candidates which might have evolved by gene duplication.

## Discussion

We explored the patterns of LD biogenesis in the ubiquitous fungal sub-phylum Pezizomycotina, which has high metabolic potential but has not yet been thoroughly analyzed. Using reverse genetics, we identified a subset of poorly characterized proteins (LDR proteins) that played a potential role in LD biogenesis. At least four LDR proteins possess a specific sequence motif, the AH, that mediates their localization to the LD surface. Particularly, LdrA is essential for regular cLD and nLD biogenesis via the AH. With bioinformatics, our study suggested gene duplication is the possible mechanism for the evolution of LDR proteins in Pezizomycotina.

Known LD-associated components are either identified with forward genetics or biochemical strategy (Guo et al, 2008; Szymanski et al, 2007; Fei et al, 2008; Currie et al, 2014; Pataki et al, 2018). However, forward genetics screening is often laborious due to genome-wide manipulation (Guo et al, 2008; Szymanski et al, 2007). Here, we applied a reverse genetics strategy for the first time by utilizing the previously established EGFP-fused library (Mamun et al, 2022 *preprint*). Colocalization analysis with LD staining dyes Nile red and BODIPY, and subsequent cross-checking with LD maker proteins Erg6 and Dga1, provided high confidence in our primary dataset consisting of 19 candidate proteins (Figs. 1–3 and Figs. S1 and 3). Multiple LDR proteins were associated as minute puncta-like structures on the ER membrane (Fig S2), leaving unanswered questions for future research, such as whether they are involved in lipid synthesis or nascent LD biogenesis.

Nearly two-thirds of LD-associated proteins had functional importance in LD biogenesis (Fig 2A–C). The reverse correlation between LD number and size observed in the individual deletion (Fig 2B and C) suggested a defect in LD growth and maturation. However, lipid profiling remains to be performed in the individual deletion strain to evaluate any alternation in lipid contents such as TG and SE. Furthermore, partially reduced colony development observed in aberrant LD-producing strains (Fig S5F) may have been caused by a somewhat inadequacy of free phospholipids for membrane proliferation due to the dominance of storage. As lipid homeostasis is crucial for growth, numerous fungicides, including azole, target it (Revie et al, 2018). CCG-8 and UsgS, the orthologues of LdrA and LdrC in *Neurospora crassa* and *Aspergillus fumigatus*, respectively are upregulated in response to azole drugs (Sun et al, 2018; da Silva et al, 2006), which provides a hypothesis that LD confers host adaptation to the antifungal drug.

The LD surface is promiscuous for interacting with AH–carrying proteins, and the binding performance is heavily influenced by amphipathic properties such as hydrophobicity, hydrophobic moment, and charge (Eisenberg et al, 1982; Prévost et al, 2018; Čopič et al, 2018; Pataki et al, 2018). AH-possessing DHRS3 with a larger axial hydrophobic surface is suggested to stably bind with LD via broad interactions with phospholipid acyl chains (Pataki et al, 2018). Here, LdrB, and LdrC contained AH with a comparatively large hydrophobic surface conferring strong LD-binding affinity to partially expel Erg6 from the LD surface (Fig S3B). Unbound (due to removal of AHs) form of LdrG was unstable in the cytosol with a similar phenomenon reported for the ubiquitous mammalian LD protein, Perilipins (Rowe et al, 2016). Many AHs are known to remain disordered in the aqueous environment and transiently folded into helical confirmation at the interface between the hydrophobic LD core and the hydrophilic cytosol (Čopič et al, 2018; Hofbauer et al, 2018). Additionally, it remains to be determined if AH-carrying LdrA, LdrB, LdrC, and LdrG are still folded in the cytosol or transiently folded at the cytosol–LD interface.

LD membrane proteome includes both anabolic and catabolic components such as lipid synthesizing enzymes and hydrolyzing lipase, respectively, and therefore plays a crucial role in LD turnover (Olzmann and Carvalho, 2019; Walther et al, 2017). For example, adipocyte differentiation-related protein (ADRP) demonstrates predominant affinity to the LD surface over adipose triglyceride lipase (ATGL), thus modulating LD dynamics (Listenberger et al, 2007). Similarly, the predominant association of LdrC resulted in a rate-limiting association of LdrA to the LD surface (Fig 7B and C), which may play a major role in the biogenesis of regular-size LD, because LdrA induced the generation supersized LD possibly via LD fusion (Fig 4E and F).

Although a metabolic switch regulated by OPi1 plays an important role in LD biogenesis in budding yeast (Loewen et al, 2004), our findings demonstrated the biophysical/structural basis of LD regulation via a novel AH of LdrA. Opi1 shows functional divergence even among yeast orthologues, such as alkane oxidation in *Yarrowia lipolytica* (Hirakawa et al, 2009) and filamentous growth in *Candida albicans* (Chen et al, 2015). Opi1 binds to the Ino2 via its activator interaction domain (AID) domain and subsequently represses genes involved in phospholipid biosynthesis (Heyken et al, 2005). The sequence similarity of Ino2 is limited within a portion of Saccharomycotina (Table S1), implying that phospholipid biosynthesis is regulated differently in Pezizomycotina. However, whether LdrA transcriptional activity (if exist any) is involved in LD biogenesis in Pezizomycotins remains to be investigated. The cellular level of LdrA is fine-tuned to regulate proper LD dynamics. The rate-limiting LdrA level (native in Fig S4A) may be owing to a bottleneck in protein folding, as previously described for Rop, a dimeric four-helix-bundle protein, where mutations at the hydrophobic core markedly increase its expression level (Munson et al, 1997) similarly to LdrA(M413N) (Fig S4A). However, the controlled LdrA level is beneficial for fungal growth, since LdrA induced lipid storage with the biogenesis of supersized cLD and nLD (Fig 4E and 5B). Supersized nLD could have a severe impact on uninucleate cells like budding yeast, which could be mitigated by limiting the nuclear import of Opi1 via primary binding with ER (Loewen et al, 2004). Alternatively, the lethal effect resulting from the nuclear degradation due to supersized nLD (Fig 5D and E) might be compensated with the multinucleate nature of *A. oryzae* in which even if some of the nuclei are disintegrated the remaining can support its survival. LdrA bound to both nLD and cLD (Fig 5A). The predominant localization of LdrA to nLD is possibly due to the absence of competitors inside the nucleus since this organelle allows only those proteins containing NLS to enter, therefore LdrA may freely associate with nLD.

Our phylogenetic analysis revealed that LDR proteins existed preferentially in Pezizomycotina, several lineages of Mucoromycota and Basidiomycota but are mostly absent or diverged in monophyletic Saccharomycotina (Fig 8 and Fig S7A-H). Yeasts undergo progressive gene loss after whole genome duplication (Kiss et al, 2019), but have evolved a specialized metabolism, known as fermentation (Novo et al, 2009; Pronk et al, 1996). Fermentation is a catabolic process that converts sugars into ethanol. Since yeast is less efficient in storing energy due to its fermentative metabolism, LDR proteins that evolved early could be lost in yeast, due to the selection pressure as irrelevant to cell physiology. On the other hand, with few exceptions, LDR proteins showed considerable existence in the several lineages of Mucoromycota (Fig 8 and Fig S7A–H). Consistently, several Mucoromycota lineages possess a remarkable potency to accumulate intracellular lipids ranging from 20 to 80% (w/w) and form lipid bodies up to 20 µm in size (Kosa et al, 2018; Dzurendova et al, 2022). Importantly, LDR proteins predominantly existed in all the major classes of Pezizomycotina and showed a tendency for gene duplication. Thus, we hypothesized that the requirement of energy storage for the adaptation to nutrient depletion in saprophytes and parasites (Naranjo-Ortiz et al, 2019) and lineage-specific metabolic versatility (Arvas et al, 2007; Wisecaver et al, 2014) could be the driving force for the evolution of LDR proteins by gene duplication to regulate lipid storage in Pezizomycotina.

## Materials and Methods

### Strains, media, growth conditions, and fungal transformation

The strains employed in this study are listed in Table S4. Transformation of *A. oryzae* was performed as previously described (Maruyama and Kitamoto, 2011). *Aspergillus oryzae* strain NSPlD1 (*niaD*^−^ *sC*^−^ Δ*pyrG* Δ*ligD*) (Mamun et al, 2020) was used as the parent strain. Using the *pyrG* selectable marker, transformants were selected using M+Met medium (0.2% NH4Cl, 0.1% (NH4)2SO4, 0.05% KCl, 0.05% NaCl, 0.1% KH2PO4, 0.05% MgSO4·7H2O, 0.002% FeSO4·7H2O, 0.15% methionine, and 2% glucose; pH 5.5). Transformants with the *niaD* and *sC* selectable markers were selected using CD medium (0.3% NaNO3, 0.2% KCl, 0.1% KH2PO4, 0.05% MgSO4·7H2O, 0.002% FeSO4·7H2O and 2% glucose; pH 5.5) and CD+Met medium supplemented with 0.0015% methionine, respectively. *A. oryzae* strains were maintained in Potato Dextrose (PD) plates (Nissui, Tokyo, Japan) at 30°C. For transformation, strains were inoculated into DPY liquid medium (0.5% yeast extract, 1% hipolypeptone, 2% dextrin, 0.5% KH2PO4, 0.05% MgSO4·7H2O, pH 5.5) and cultured overnight with constant shaking. For microscopy, conidia were inoculated on glass base dishes containing 100 μL CD liquid medium supplemented with 1% casamino acids.

### Plasmid construction and strain development

#### Construction of strains expressing mCherry-tagged LD-associated proteins

The open reading frames of 19 LD-associated proteins were amplified from RIB40 genomic DNA with PrimeSTAR^®^ HS DNA Polymerase (TaKaRa, Otsu, Japan) for high-fidelity PCR, using primers listed in Table S5. The amplified fragments were then fused with *Sma*I-linearized pisCIIA-mCherry (Mamun et al, 2020) using the In-Fusion HD Cloning Kit (Clontech Laboratories, Mountain View, CA, USA). Recombinant plasmids for the expression of mCherry-fused 19 proteins were digested with *Not*I and ectopically inserted into the ATP sulfurylase (*sC*) locus of the wild-type strain NSlD1 (Mamun et al, 2020) by homologous recombination (Fig S1B and C).

#### Generating LDR gene deletions library

Primers used for the generation of the deletion strains (Fig 2A-C) are listed in Table S5. Replacement of individual genes with the Orotidine-5’-phosphate decarboxylase (*pyrG*) marker was performed by amplifying 1.5 kb upstream and 1.5 kb downstream of the gene of interest using the primer sets gene-upstream1_F/gene-upstream2_R and gene-downstream1_F/gene-downstream2_R, respectively. The *pyrG* marker was amplified from RIB40 genomic DNA using the primers PyrG_F and PyrG_R. The three amplified DNA fragments (upstream, and downstream regions of targeted genes and *pyrG*) and linearized-pUC19 vector were fused using the In-Fusion HD Cloning Kit. Deletion constructs were further amplified using the primer set gene-upstream1_F/gene-downstream2_R from the resultant plasmid, and then introduced into the native locus of the *A. oryzae* strain NSPlD1(Mamun et al, 2020) by homologous recombination. Transformants were selected using uridine/uracil prototrophy.

#### Generating double deletion straains

To generate double deletions (Figs 2A–C and 7A), 1.5 kb upstream and 1.5 kb downstream of *ldrF*, *ldrH*, and *ldrA,* were amplified from RIB40 genomic DNA using the primer sets gene-upstream1_F/gene-upstream2_R and gene-downstream1_F/gene-downstream2_R, respectively listed in Table S5. The *sC* marker was amplified separately. The three amplified DNA fragments (upstream, and downstream regions of targeted genes and *sC*) and linearized-pUC19 vector were fused. Deletion constructs were further amplified using the primer set gene-upstream1_F/gene-downstream2_R from the resultant plasmid. PCR-amplified deletion constructs of *ldrF*, *ldrH*, and *ldrA* were introduced into the native loci of Δ*ldrE,* Δ*ldrG,* and Δ*ldrC,* respectively.

#### Expression of EGFP-tagged LDR and SPP proteins from the endogenous promoter

For complementation (Fig 2D and E), the DNA fragments from 1.4 kb upstream until the stop codon and 0.6 kb downstream of the respective genes were amplified separately from RIB40 genomic DNA using primers listed in Table S5. EGFP sequence was amplified from pUt-EGFP (Mamun et al, 2020) using EGFP_F and EGFP_R. The amplified three fragments (upstream + ORF, downstream, and EGFP) were fused with a *Sma*I-linearized pUXN (Mamun et al, 2020) vector, yielding pU*gene*N-EGFP. The resultant plasmids were digested with *Not*I and introduced into the *niaD* locus of the corresponding deletion strain by homologous recombination. For negative control, *Not*I-digested PUXN vector was introduced into the *niaD* locus of the individual deletion strain.

#### Construction of strains expressing mCherry-tagged Aoerg6 and AoDga1 for colocalization with EGFP-tagged LDR proteins

For colocalization analysis (Fig 3 and Fig S2 and S3), *A. oryzae* proteins AO090026000845, AO090011000289, and AO090005001238 were identified as orthologues of Dga1, Erg6, and Sec63, respectively using budding yeast proteins. The open reading frame (ORF) of three respective genes were amplified from RIB40 genomic DNA using primers listed in Table S5. The amplified fragments were then fused with *Sma*I-linearized pisCIIA-mCherry (Mamun et al, 2020). Recombinant plasmids expressing mCherry-fused proteins were digested with *Not*I and ectopically inserted into the *sC* locus of the NSlD1. The resultant strains expressing AoDga1-mCherry and AoErg6-mCherry were further transformed (ectopically) with *Not*I digested EGFP-fused plasmids (Mamun et al, 2022 *preprint*) into the *niaD* locus by homologous recombination. Strain expressing AoSec63-mCherry was transformed with 12 EGFP-fused LD-associated proteins (Fig S2).

#### Construction of truncations and AH-disrupting mutants of LDR proteins

To truncate the putative AHs (Figs 4 and 6), the flanking ORF fragments before and after AH were separately amplified from the corresponding EGFP-fused plasmids bearing full-length sequence. To construct AH disrupting mutants, the customized primers containing 15 bp overlapping sequences, which bear the codon of desired mutations were used to amplify the flanking ORF fragment before and after the targeted sites. To develop double such as LdrC(V175N/V168N), flanking ORF fragments before and after the targeted sites were amplified from the corresponding formerly developed single mutation-bearing plasmid such as LdrC(V175N). Three fragments including upstream flanking, downstream flanking, and *Sma*I-linearized either pUt-EGFP (LdrA variants in Fig 4C; LdrG variants in Fig 6I) or pisCIIA-mCherry (LdrC, and LdrB variants in Fig 6C, F) were fused. The resultant plasmids were digested with *Not*I and then introduced either into the *niaD* (LdrA, and LdrG) or sC (LdrC, and LdrB) locus of the corresponding deletion strain by homologous recombination. At the same time, plasmids corresponding to wild-type sequences were also inserted into the respective deletion background for later use as the positive control.

#### Expression of NLS (AoStuA)-, AoNup84-, and AoNup49-mCherry

To visualize the nuclear periphery (Figs. 5 and 6), components of the nuclear pore complex, Nup84 and Nup49 were used. *Aspergillus oryzae* proteins AO090138000215 and AO090023000199 were identified as the orthologues of *Aspergillus nidulans* corresponding proteins (Liu et al, 2009). The ORF of AoNup84 and AoNup49 were amplified from RIB40 genomic DNA using primers listed in Table S5. Similarly, to visualize the nucleus or nucleoplasm (Fig 5), the C-terminal region of StuA, referred to as NLS (400–645aa, which contains NLS located outside of the functional domain) was amplified from RIB40 genomic DNA. All the amplified fragments including AoNup84, AoNup49, and NLS were then fused individually with *Sma*I-linearized pisCIIA-mCherry. Recombinant plasmids were digested with *Not*I and ectopically inserted into the *sC* locus of the Δ*ldrA* and wild type. The aforementioned plasmids containing EGFP-fused mutants and truncated variants of LdrA were digested with *Not*I and then introduced either into the *niaD* of Δ*ldrA* expressing AoNup84-mCherry. Here, for the development of the untagged (EGFP) version (Figs. 5D-E; 6), the DNA fragments of LdrA variants including truncated, mutated, and wild-type were amplified until the stop codon from the above developed (Fig 4C) EGFP-fused respective plasmids, and subsequently fused with *Sma*I linearized pUtNAN (Mamun et al, 2020). To construct LdrA variants expressed from the endogenous promoter, the fragments 1.4 kb upstream and those from the start codon till T*amyB* were separately amplified from RIB40 genomic DNA and the corresponding plasmids (aforementioned as untagged version), respectively. The two amplified fragments (upstream including promoter and ORF + T*amyB* terminator) were fused with the *Sma*I-linearized pUXN vector. Resultant plasmids bearing LdrA variants expressed from the heterologous and endogenous promoter were digested with *Not*I and then introduced into the *niaD* locus of Δ*ldrA* expressing NLS (StuA)-mCherry (Figs. 4-5) and AoNup49-mCherry (M413N in Fig 5).

#### Co-expression of LdrA-EGFP and LdrC-mCherry, as well as overexpression of LdrA in ΔldrAΔldrC

For co-expression (Fig 7B-C), a full-length LdrA fragment was amplified from RIB40 genomic DNA using primers listed in Table S5. The amplified fragment was then inserted into the *Sma*I site of pUtNAN (Mamun et al, 2020), yielding pUtNA*ldrA*N. The subsequent plasmid and aforementioned pUt*ldrA*-EGFP were digested with *Not*I and ectopically introduced into the *niaD* locus of NlD1-LdrCmChry and NlD1-LdrC(V175N/V168N)mChry. For complementation (Fig 7A), *Not*I digested pUtNA*ldrA*N was introduced into the *niaD* of double deletion *ldrAldrC* using homologous recombination.

### Fluorescent microscopy and image processing

Living hyphae expressing EGFP and mCherry-tagged proteins or staining with LD dye BODIPY and Nile red were observed using an IX71 inverted frame confocal microscope (Olympus, Tokyo, Japan) equipped with 100× Neofluar objective lenses (1.40 numerical aperture); 488-(Furukawa Electric, Tokyo, Japan) (BODIPY, EGFP) and 561-nm (Melles Griot, Rochester, NY, USA) (Nile red, mCherry) semiconductor lasers, GFP filters (Nippon Roper, Tokyo, Japan); a CSU22 confocal scanning system (Yokogawa Electronics, Tokyo, Japan), and an Andor iXon cooled digital CCD camera (Andor Technology PLC, Belfast, UK). Images were analyzed using Andor iQ 1.9 software (Andor Technology PLC) and exported as TIFF files.

To analyze the percentage of AoErg6-mCherry and AoDga1-mCherry colocalized with EGFP-fused LDR proteins, the total mCherry-(population 1) and EGFP-positive (population 2) area regardless of the particular distribution pattern was calculated using the color threshold tool in ImageJ, version 15.3 (Schneider et al, 2012). The area corresponding to the colocalized intersection between EGFP and mCherry channels was calculated by adjusting a similar set of the color threshold tool (population 3). Quantification settings were identical throughout the experiment. The Percentage of colocalization for mCherry channel was calculated using the following formula (Rowe et al, 2016): Population 3/ Population 1 × 100 (Rowe et al, 2016). The Percentage of colocalization for EGFP channel was calculated using the following formula: Population 3/ Population 2 × 100.

The colocalizations of BODIPY- or Nile red-stained LD with mCherry- or EGFP-tagged LDR proteins, respectively were analyzed using FIJI. The total number of LDs and the colocalized LD with LDR proteins were counted using the particle detection tools of Plugins of FIJI. The colocalization percentage was calculated from colocalized-LD divided by the total number of LD and multiplied by 100 (Colocalization percentage = colocalized-LD/ total LD X 100).

Similarly, the nLD number (Fig 5) was counted using the colocalization of NLS-mCherry positive with BODIPY-stained nLD from the hyphal area except for the apical compartment. The total number of nuclei was then counted with the specified area and the value was presented as a percentage.

To quantify fluorescent intensity, the images from EGFP or mCherry-fused LDR variants were captured at identical excitation and detection settings from live hyphae. Fluorescent intensity was measured by drawing the single transverse line on the LD membrane (Fig. S4E and S6D and E) or the nucleus (Fig S4C) or across hyphae (Fig 5E) using the plot profile tool in ImageJ (Schneider et al, 2012).

### LD staining and quantification

Strains were cultured on the liquid minimal medium in glass bottom dish for 20 h, and then medium was replaced with fresh medium containing either 0.5 μg/ml BODYPY 493/503 (Sigma, St. Louis, Missouri, USA) or 10 μg/mL Nile red (Sigma, St. Louis, Missouri, USA). The culture containing the staining solution was incubated for 10 min at room temperature. Due to the filamentous shape of fungal hyphae, the hyphal areas covering one window size (with ×100 lens) equivalent to 45 µm were captured preferentially from a uniform and non-clustered LD region. Fluorescent images were taken with ×100 objective lens and captured with an 8-bit setting and subsequently stored in the tiff file format. The images captured from at least three independent experiments were analyzed using ImageJ software. LD number and size within the analyzed hyphal area were quantified using the “analyze particles” tool in ImageJ with size (pixel^2^) settings from 0.1 to 1000 and circularity from 0 to 1. Identical thresholds were applied to all experimental conditions. The values were presented as average.

### Western blotting

Strains were cultured on minimum medium (CD liquid medium) for 20 h to induce LD biogenesis. The fungal mycelium was harvested using liquid nitrogen and subsequently grinded with multi bead shocker. After extracting, cell lysates were incubated with the buffer containing detergent NP-40 (50 mM Tris-HCl (pH 8.0), 200 mM NaCl, 1 mM/ mL PMSF, 1 mM/ mL Protease inhibitor, 1% NP-40) to solubilize the membrane protein. Mycelial extract solubilized with the buffer was incubated on ice for 20 min. After centrifugation with 1000x *g* for 10 min, the supernatant was collected. The protein samples were separated using 10% SDS-PAGE, and then transferred to Immobilon-P polyvinylidene difluoride (PVDF) membranes (Millipore, Bedford, MA, USA) using a semidry blotting system (Nihon Eido, Tokyo, Japan). To detect EGFP, Living Colors^®^ A.V. monoclonal antibody (cat # 632380, 1: 2,000, Clontech) and horseradish peroxidase (HRP)-labeled anti-mouse IgG (H + L) (cat #, 7076S, 1: 2,000, Cell Signaling Technology, Danvers, MA, USA) were used as primary and secondary antibodies, respectively (Fig 5G). Chemiluminescence was detected using a Western Lightning-ECL system (PerkinElmer, Waltham, MA, USA) and an LAS-4000 image analyzer (GE Healthcare, Buckinghamshire, UK).

### Prediction of Amphipathic helices

To predict the AH, all the LDR proteins were comprehensively analyzed with protein fold recognition server, Phyre2 (Kelley et al, 2015) to retrieve the α-helix region. Each α-helix was then analyzed with HELIQUEST (Gautier et al, 2008) to know the helical wheel orientation. If the helical wheel showed segregation of helix into hydrophobic face with considerable length and polar face especially enriched with positively charged residues, then these are considered. The α-helix with hydrophobicity (<H>) and hydrophobic moment (<µH>) around 0.35 were set as the criteria to be taken (Čopič et al, 2018). The predicted AH was further analyzed with AlphaFold2 (Mirdita et al, 2022) and Chimera 1.16 to evaluate the distribution of hydrophobic and polar residues in the helix.

### Phylogenetic analysis

LDR proteins were investigated using Simple Modular Architecture Research Tools (SMART: http://smart.embl-heide lberg.de/) (Schultz et al, 1998) to identify predicted domains. The domain organizations were further cross-checked with NCBI and InterPro site. Multiple sequence alignments were performed to analyze sequence conservation using ClustalW (Thompson et al, 2002).

We followed a previously described method for analyzing substitution rate (Fig 8) of each LDR protein from the Pezizomycotina root (Nguyen et al, 2017). Briefly, we performed BLASTp (≤ 1.0e-5) to search and subsequently retrieve the best hit protein from representative fungal species (Table S3A) using individual LDR proteins as the query. Multiple sequence alignments were built for individual proteins using ClustalW (Thompson et al, 2002) with default setting of gap penalties. Maximum likelihood trees were generated for each LDR protein using PhyML. Phylogenetic trees were crosschecked by further constructing them with molecular evolutionary genetics analysis (MEGA) tool (Kumar et al, 2016). The substitution rate (Table S3C) was calculated for an individual species x in the corresponding tree, the evolutionary distance between protein sequences and Pezizomycotina ancestral sequences of *A. oryzae* LDR proteins was estimated by the score: 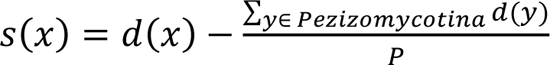 (Unit: substitutions/site), where *d*(*x*) indicates the branch length from species *x* to the Pezizomycotina root. *P* represents the number of species *y* in Pezizomycotina used in phylogenetic analysis, and therefore 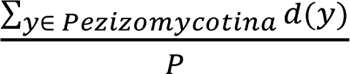 represents the average branch length from each species *y* in Pezizomycotina to the Pezizomycotina root.

Reciprocal BLAST (Hernández-Salmerón et al, 2020; Li et al, 2003; Ward et al, 2014) was employed to find orthologues in *A. oryzae* (Table S3B) with the best hit candidates (Table S3A), used in substitution rate analysis in Fig 8) as the query. To elucidate the pattern of gene evolution, additional phylogenies were generated using fungal proteins with shared domains (Fig S7A-H). Total proteins within a given species containing analyzed domains were retrieved from the FungiDB genome database. Domain information for those species absent in the FungiDB database was retrieved from UniProt. Phylogenetic trees for analyzed proteins were constructed using Mega X (Kumar et al, 2016).

### Statistical analysis

The results of at least three independent experiments are presented as means. Error bars represent standard deviations as indicated in the figure legends. Statistical significance was tested with the two-tailed Student’s *t-*test using Microsoft Excel; significance is indicated as ns-not significant *p* > 0.05, \**p* < 0.05, \*\**p* < 0.01, \*\*\**p* < 0.001, \*\*\*\**p* < 0.0001

## Acknowledgments

We are highly grateful to Professor Jun-Ichi Maruyama, Brewing Microbiology Lab, department of Biotechnology, The University of Tokyo, Japan, for laboratory support, including availability of all the reagents and instruments used in this study from his own research fund (KAKENHI Grant Number 21H02098). We are also grateful to Dr. Shinichi Nishimura Microbiology Lab, The University of Tokyo for providing BODIPY 493/503 and related suggestions for the study.

## Author contributions

M. A. A. Mamun conceived of and designed the study. M. A. A. Mamun performed all the experiments and interpret the results. M. A. A. Mamun, M. A. Reza, and M. S. Islam performed the bioinformatics analysis. Reza and M. S. Islam provided relevant scientific advice to conduct the study. M. A. A. Mamun, M. A. Reza, and M. S. Islam wrote and approved the manuscript.

## Data availability

All data supporting the findings of the present study are available in this article and Supplementary Information files, or from the corresponding authors upon request.

## Abbreviations

LD: lipid droplets

LDR: lipid <u>d</u>roplet regulating

LD: Cytoplasmic

nLD: cLD Nuclear

AH: amphipathic helices

NLS: nuclear localizing signal

HRP: horseradish peroxidase

## Footnotes

* The manuscript is now under revisions (third) in *Nature communications* (NCOMMS-22-01369C).

## Online supplemental material

Figure S1 provides additional information about the colocalization of novel LD-associated proteins with neutral lipid dyes (Supporting Figure 1). Figure S2 provides information about the colocalization of novel LD-associated proteins with ER network (Supporting Figure 1). Figure S3 provides information related to the colocalization of several LDR proteins with LD marker proteins Erg6 and Dga1 (Supporting Figure 3). Figure S4 presents information related to AH-mediated localization of LdrA to the LD membrane (Supporting Figure 4). Figure S5 provides information on the importance of AH to bind to and regulate LD biogenesis (Supporting Figure 4 and 5). Fig S6 provides information related to the putative AHs of LdrC, LdrB, and LdrG (Supporting Figure 6). Figure S7 presents phylogenetic trees which validate the relationship between orthologues and gene duplication of the LDR proteins (Supporting Figure 8).

---

**Figure S1.**
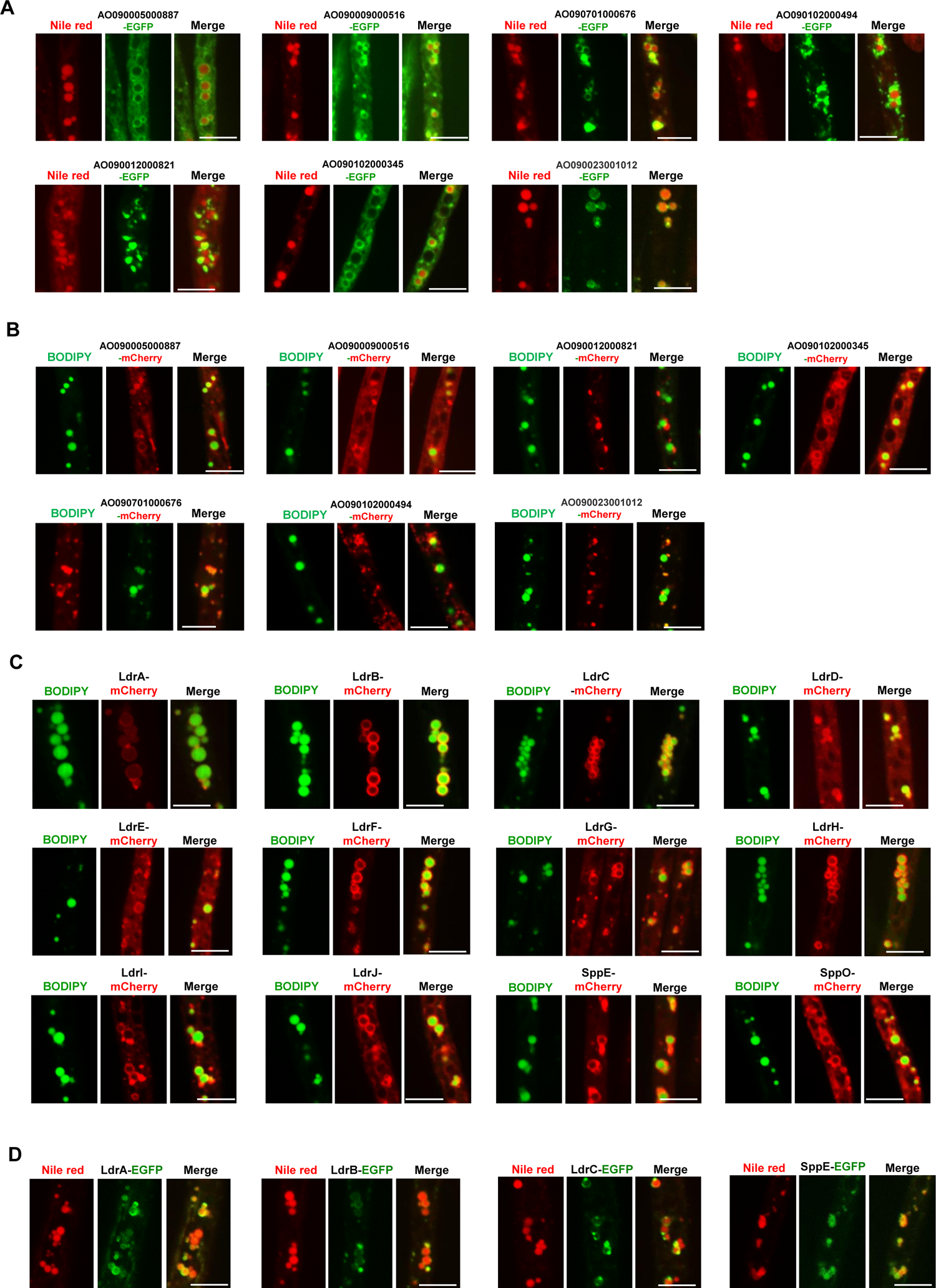
Colocalization of LD-associated proteins with LD marker dyes. Strains were grown on the minimum medium for 20h. Representative microphotographs were presented. **(A)** Colocalization of LD-associated EGFP-fused proteins (functionally not important) with Nile red was observed from living hyphae. **(B)** Colocalization of LD-associated mCherry-fused proteins (functionally not important) with BODIPY. **(C)** Colocalization of mCherry-fused LDR and two SPP proteins with BODIPY. **(D)** Colocalization of LD-associated EGFP-fused proteins expressed from the endogenous promoter with Nile red. **(A, D)** LDs were visualized with 10 µg/ mL Nile red. **(B, C)** Fungal hyphae were stained with 5µM BODIPY for the visualization of LD. Scale bars 5 µm.

**Figure S2.**
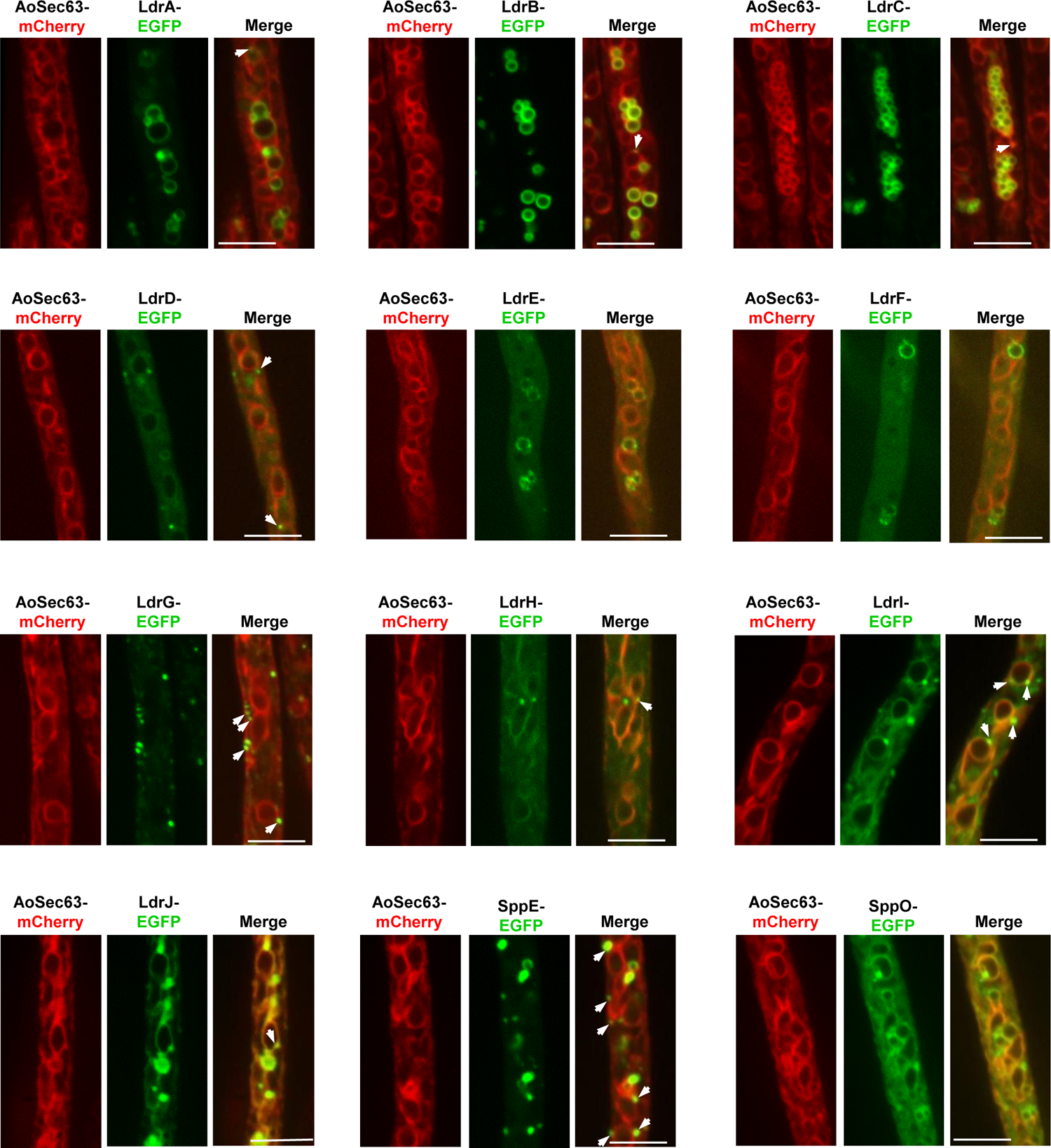
Colocalization of LDR- and two SPP proteins with ER marker Sec63. Strains were grown on the minimum medium for 20h. Representative microphotographs were presented. ER network was visualized with ER marker protein Sec63 fused with mCherry. EGFP-fused LDR and two SPP proteins expressed from *amyB* promoters were introduced into the stains expressing AoSec63-mCherry. Arrowheads indicate dot-like localization on the ER membrane. Scale bars, 5 μm.

**Figure S3.**
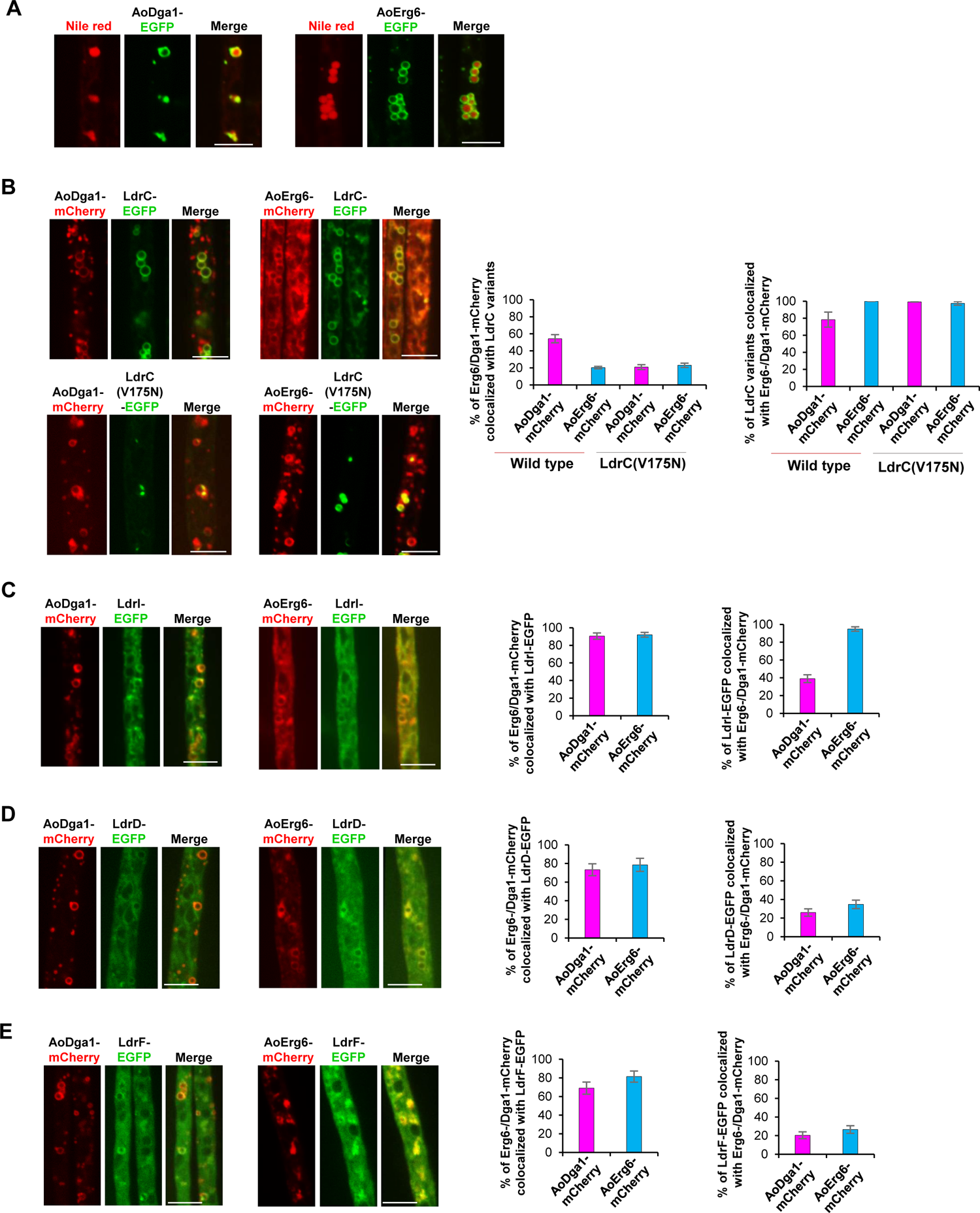
Colocalization of LDR proteins with LD marker proteins Erg6 and Dga1. **(A)** Colocalization of AoErg6-EGFP and AoDga1-EGFP with LD marker Nile red. **(B-E)** AoErg6- and AoDga1-mCherry under the control of *amyB* promoter were introduced individually into wild-type. EGFP-tagged LdrC, LdrD, LdrF, and LdrI from *amyB* promoter were then introduced into the resultant strains. Strains were cultured on the minimal medium with optimized carbon concentration. The percentage of AoErg6-and AoDga1-mCherry associated with EGFP-tagged analyzed LDR proteins and *vice versa* were calculated using ImageJ. Data are shown as mean ± SD, n = 12 hyphae. **(B)** Colocalization of LdrC- and LdrC(V175N)-EGFP with AoErg6-and AoDga1-mCherry. **(C)**, **(D)**, and **(E)** Colocalization of AoErg6- and AoDga1-mCherry individually with LdrI-, LdrD-, and LdrF-EGFP, respectively. Bars 5 µm.

**Figure S4.**
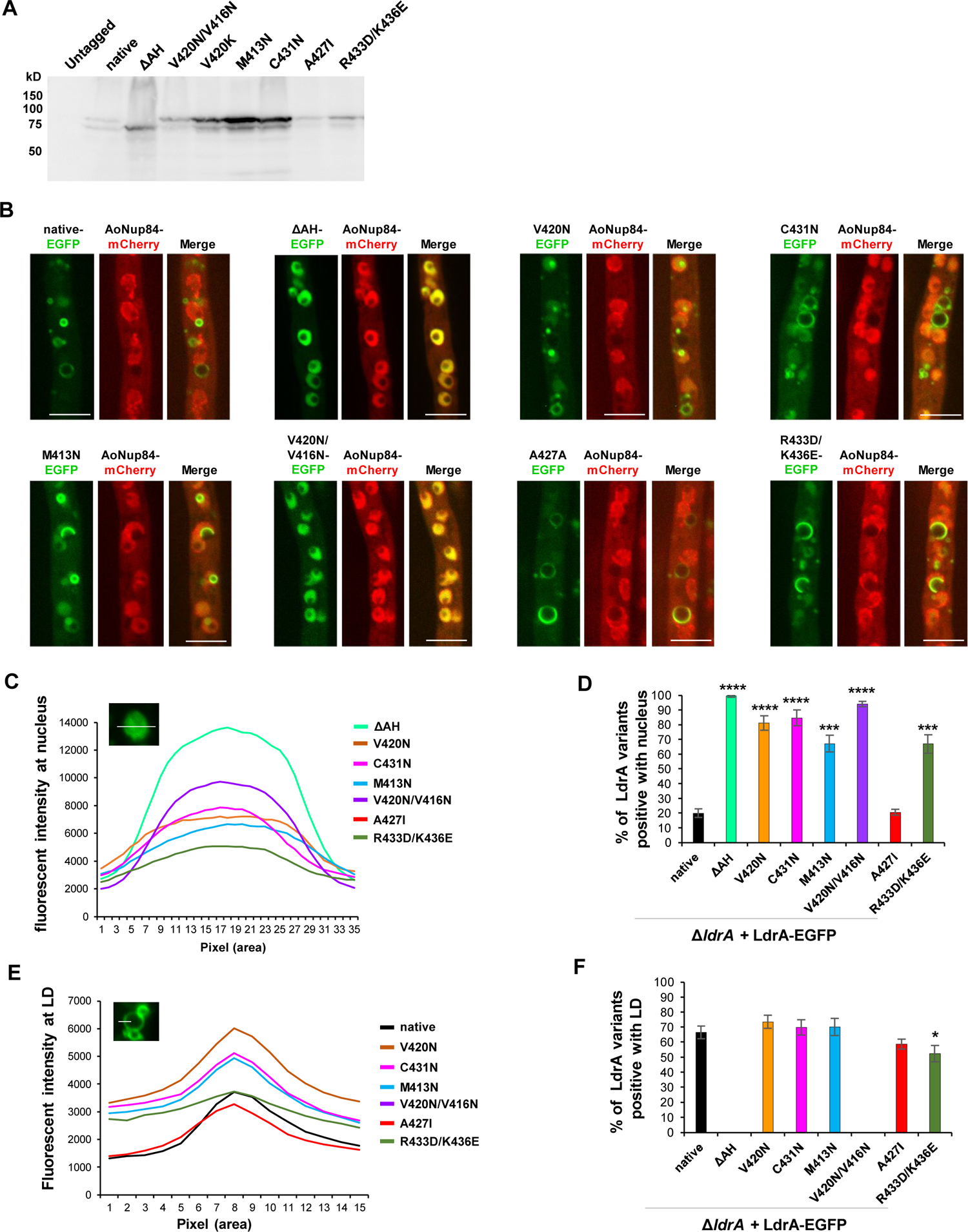
Putative AH of LdrA modulates its subcellular localization. EGFP-tagged AH disruptive-mutants expressed from the *amyB* promoter were introduced into Δ*ldrA.* **(A)** Expression and size of the variants was detected by western blotting using anti-GFP antibody. **(B-D)** For nuclear enrichment analysis, Δ*ldrA* ectopically expressing AoNup84-mCherry was introduced with EGFP-fused LdrA variants. **(B)** Colocalization of EGFP-fused LdrA variants with AoNup84-mCherry. Scale bars 5µm. **(C)** Nuclear fluorescent intensities of LdrA variants were calculated using ImageJ. n = 30 nuclei. **(D)** Percentage of AoNup84-positive nuclei with LdrA variants. n ≥ 212 Nup84-marked nuclei. **(E)** Fluorescent intensity at LD membrane was calculated using ImageJ. n = 30 LDs. **(F)** After grown the EGFP-tagged variants in CD medium, LDs were visualized with 10 µg/mL Nile red. Percentage of Nile red-stained LD positive with LdrA variants. n ≥ 221 LDs. **(C, F)** The bar diagrams are represented as the mean value from three independent experiments. Statistical significance was assessed using two-tailed student *t-test*. Error bars represent SD. ns-not significant p > 0.05, *p < 0.05, **p < 0.01, ***p < 0.001, ****p < 0.0001.

**Figure S5.**
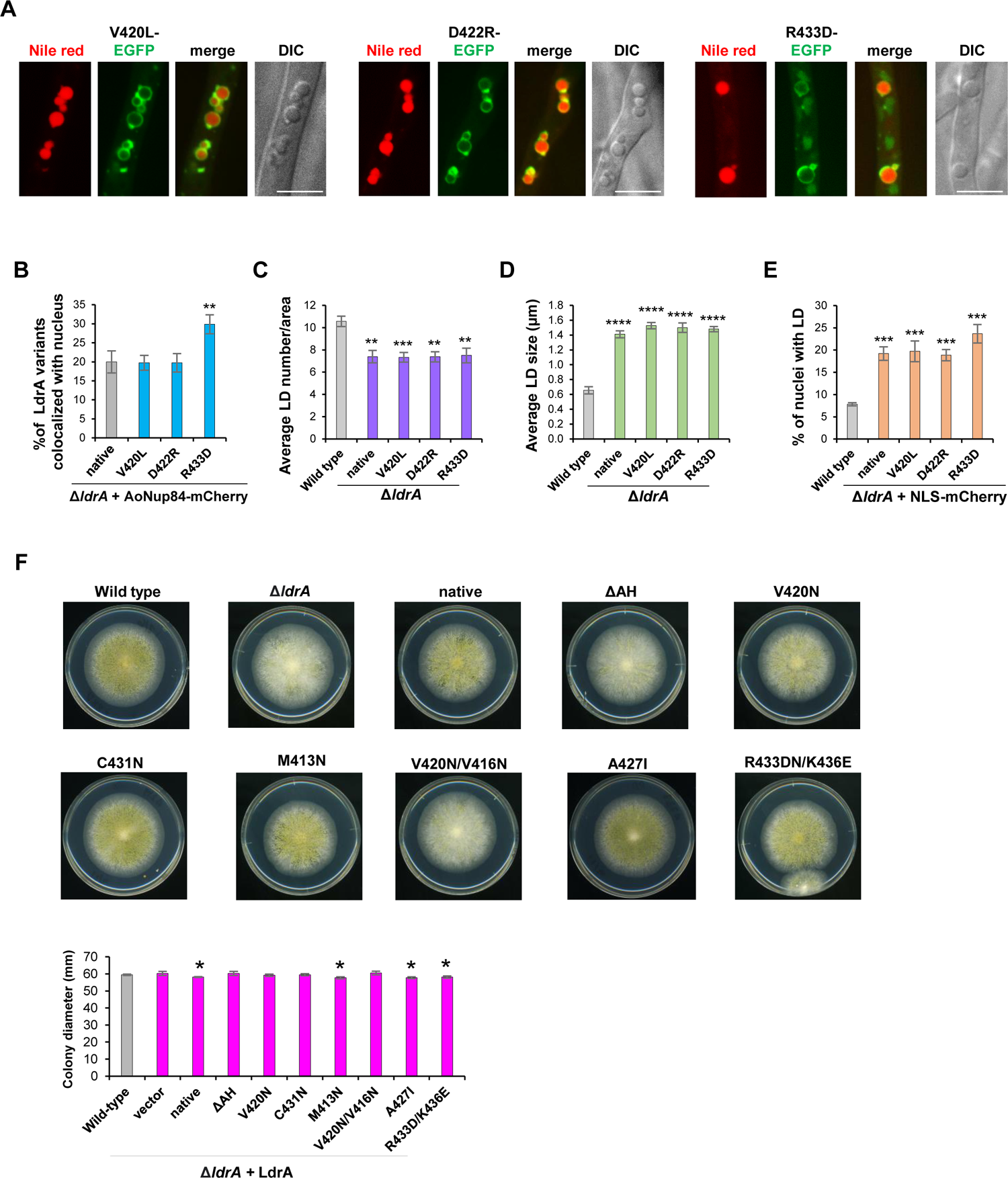
Importance of the putative AH of LdrA to target the LD surface. **(A)** EGFP-tagged LdrA variants having mutations in the putative AH were introduced into Δ*ldrA* from the *amyB* promoter. Colocalization of LdrA variants with LD marker Nile red. Scale bars 5 µm. **(B)** Percentage of ApNup84-mChrry-labelled nucleus positive with LdrA variants. n ≥ 212 nuclei **(C-D)** Quantification of LD number (per area) and LD size from LdrA variants (no EGFP tagging), respectively. **(C)** n ≥ 21 hyphal area and **(D)** n ≥133 LD. **(E)** Percentage of nuclei positive with LDs. n ≥ 160 nuclei. **(F)** A partially reduced conidiation and colony growth in strains with defective and overactive in LD biogenesis, respectively. A conidial suspension (10^4^ conidia in 4 µl water solution) of the strains was spotted on PD media and cultured for 4 d. **(B-F)** Bar diagrams are presented as the mean value from three independent experiments. Statistical significance was assessed using the two-tailed student *t-test*. Error bars represent SD. ns-not significant p > 0.05, *p < 0.05, **p < 0.01, ***p < 0.001, ****p < 0.0001

**Figure S6.**
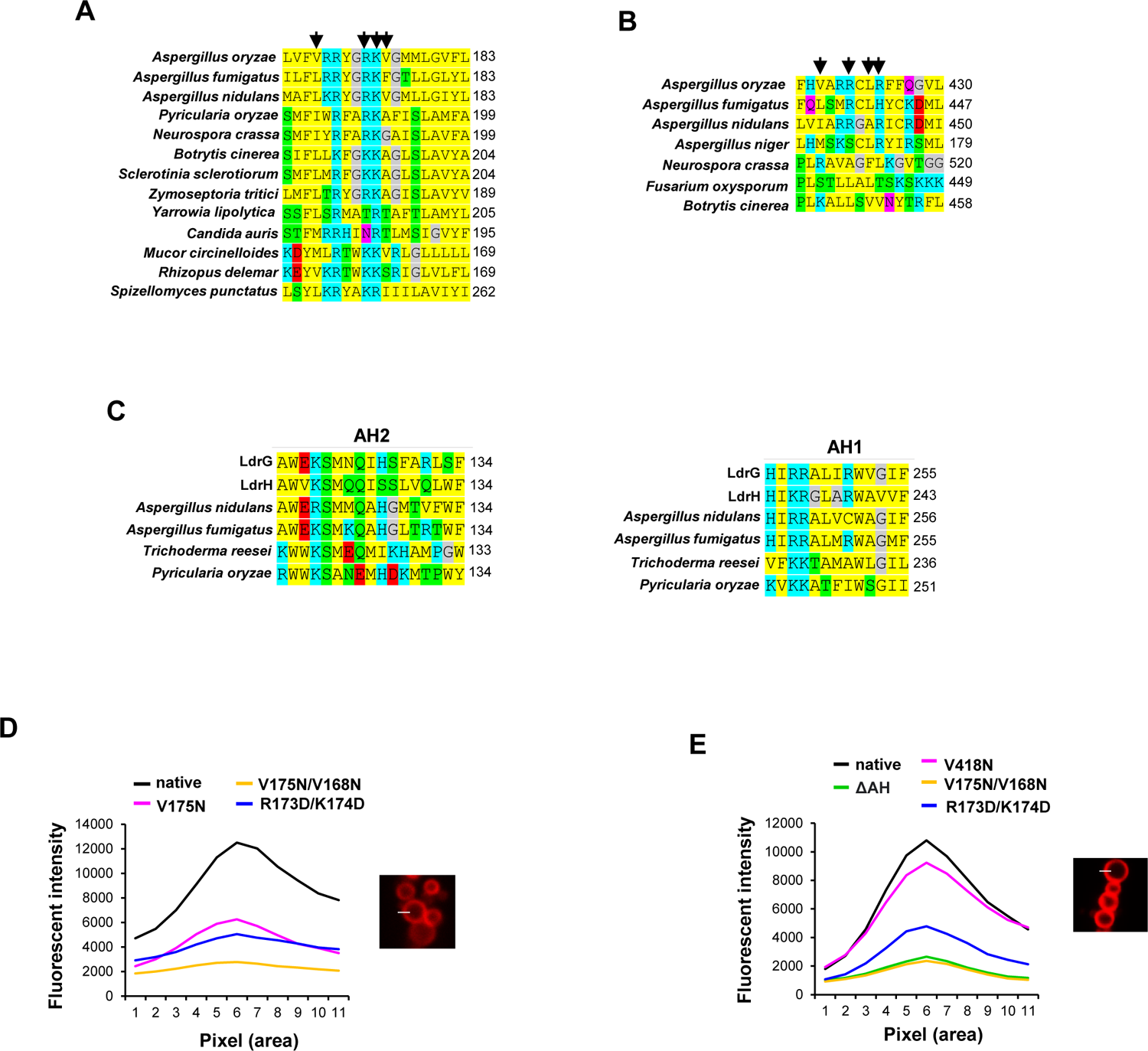
Analysis of the predicted amphipathic helices of LdrC, LdrB, and LdrG. Multiple sequence alignment of AH regions using ClustalW. The sequences were retrieved from representative fungal orthologues. Amino acid properties are indicated by colors. **(A)** LdrC, **(B)** LdrB and **(C)** LdrG. (B, C) arrows denote the residues that were substituted for AH-disruption. Mean fluorescent intensity of the mCherry-fused variants of LdrC **(D)**, and LdrB **(E)**. Fluorescent intensity was measured by drawing line (covering 15 pixels) on the LD membrane and flanking regions using ImageJ. n = 20 LDs.

**Figure S7.**
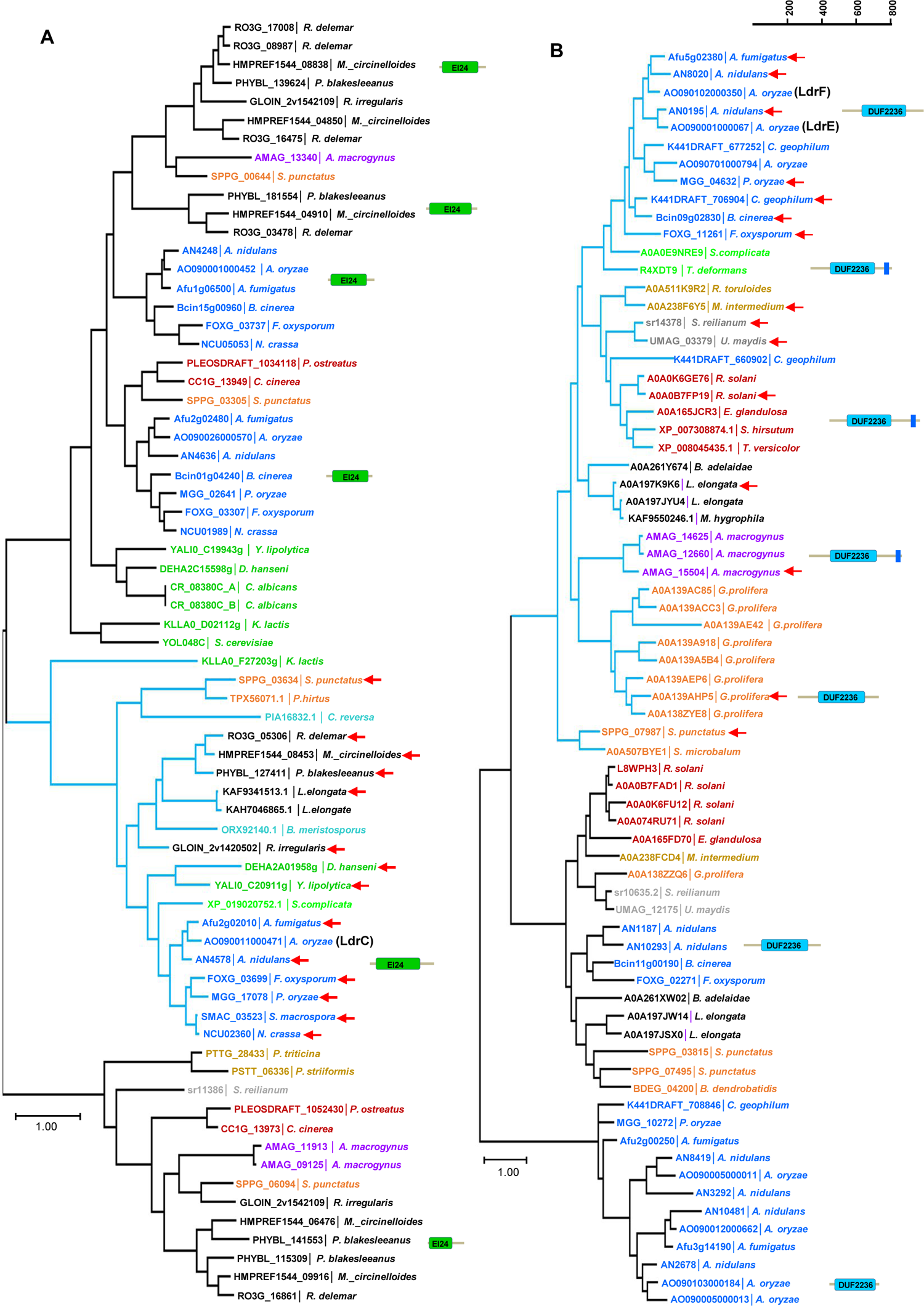
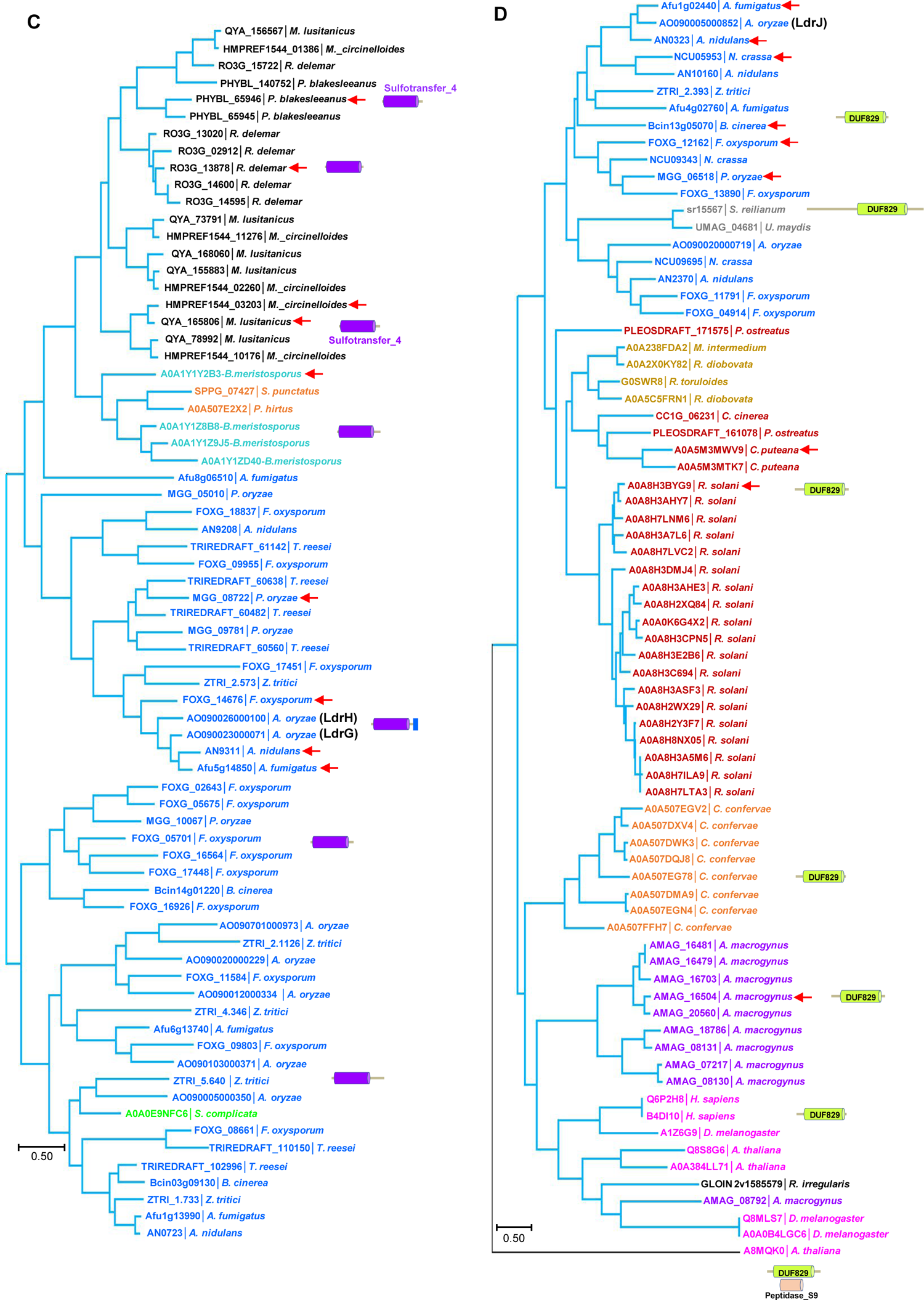
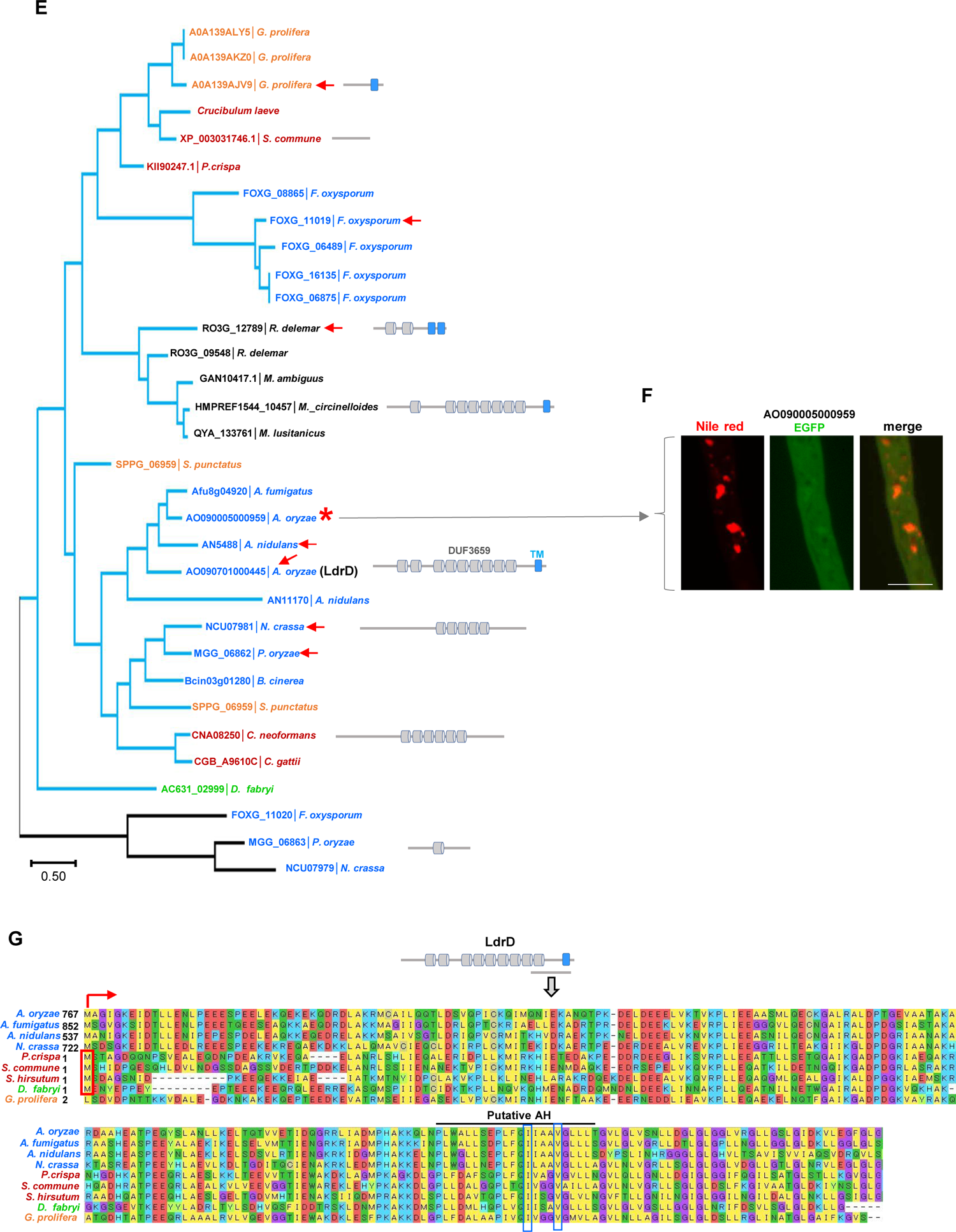
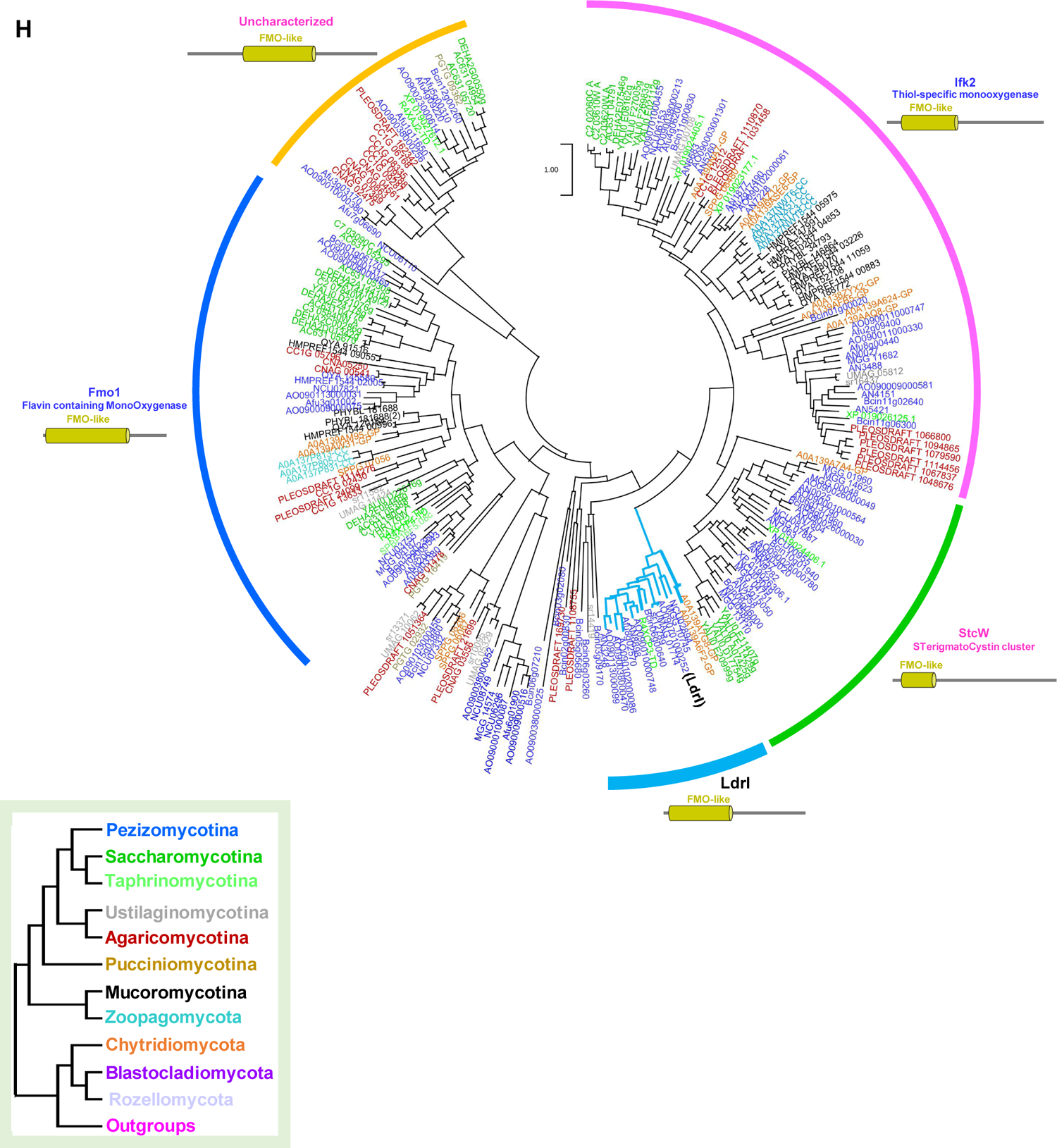
Phylogenetic trees of eight LDR proteins. Domains were predicted using the simple modular architecture research tool (SMART) and cross-checked in NCBI. Total proteins containing the analyzed domain within the given species were retrieved from the FungiDB genome database and case dependently from UniProt (See Methods). Maximum-likelihood trees were constructed using Mega 11. Diagrams of protein domain structures are shown with the respective clades. The possible orthologous clades are represented by the blue color branches of the tree. In the cases of multiple proteins with the same domain, red arrowheads indicate the proteins used for the substitution rate analysis in Fig. 8. **(A)** Phylogeny of fungal proteins having EI24 present in LdrC. **(B)** Phylogeny of fungal proteins having DUF2236 domain present in LdrE and LdrF. **(C)** Phylogeny of fungal proteins having Sulfotransfer_4 domains present in LdrG and LdrH. **(D)** Phylogeny of fungal proteins containing DUF829 domains present in LdrJ. **(E)** Phylogeny of fungal proteins having DUF3659 domains in LdrD. Star mark represents the paralogues of LdrD. **(F)** AO090005000959 fused with EGFP was ectopically expressed into the wild-type. LD was visualized with Nile red. Bars 5 µm. **(G)** Multiple sequence alignment shows the conserved C-terminal region presenting the critical residues for amphipathic helices organization within boxes. **(H)** Phylogeny of fungal proteins containing FMO-like domains present in LdrI. Gene names ending in CC, GP, and TD indicate *Conidiobolus coronatus, Gonapodya prolifera,* and *Taphrina deformans*, respectively.

